# Physical exercise and brain network dynamics: reduction of frontoparietal-striatal connectivity following 1 hour of aerobic cycling

**DOI:** 10.1101/2025.08.28.672547

**Authors:** Jetro J. Tuulari, Wajiha Bano, Tiina Saanijoki, Joana Cabral, Henrique Fernandes, Marina Charquero Ballester, Jakub Vohryzek, Lauri Tuominen, Lauri Nummenmaa, Kari Kalliokoski, Morten L. Kringelbach, Jussi Hirvonen

## Abstract

Physical exercise is beneficial for metabolic health and cognitive performance. In addition, exercise can be highly pleasurable and and serves as an effective stress reliever. Extant studies have highlighted frontal, parietal, and subcortical brain changes following acute bouts of exercise. However, slower, slightly delayed brain correlates after exercise at the network level have not been studied. Therefore, this study’s objective was to investigate the changes in dynamic functional connectivity after 60 minutes of exercise in healthy males.

Here we measured a 6-minute resting state fMRI in 24 young males at baseline and after a 60-minute cycling exercise challenge. Apart from routine preprocessing, the data were denoised with FSL-FIX and modeled with i) leading eigenvector dynamics analysis (LEiDA) to probe whole brain network dynamics and ii) with group independent component analysis (ICA) and dual regression to quantify static brain connectivity. The within subject statistical tests compared baseline to post-exercise conditions.

We found that a striato-fronto-parietal network is destabilized after exercise, as indicated by a lower probability of occurrence in dynamic analysis through LEiDA. The brain areas in the network include the bilateral caudate, putamen and pallidum as well as middle orbitofrontal, frontal operculum, frontal trigonum and inferior parietal cortices. No differences between baseline and post exercise conditions were found in the dual regression of the group ICA components.

We conclude that 60 minutes of cycling causes a prolonged effect in brain network dynamics, reducing synchronization between the striatum and frontoparietal networks with respect to baseline. This provides insights into the network-level neural correlates of aerobic exercise, which may be directly linked with the stress relieving effects of physical exercise.

## INTRODUCTION

Physical exercise benefits metabolic and mental health (Myers et al., 2019). Apart from the well-established general health-promoting effects, exercise also positively affects cognitive performance (Raichlen and Alexander, 2017). Physical activity is rewarding, and endorphins are likely mediators of the exercise-induced pleasurable feelings (Nummenmaa et al., 2018; Saanijoki et al., 2018a, 2018b). Exercise, across varying degrees and protocols, improves cognitive task performance, elevates mood, and decreases stress (Basso and Suzuki, 2017). Cerebral effects of exercise are thus of interest as possible mediators of increased well-being and brain health.

One prominent hypothesis on the acute effects of exercise is the hypoactivation of the (pre)frontal cortical areas, which are involved in cognitive processing as compared to areas that are required for motor function, such as the primary motor cortex, striatum, and cerebellum (Dietrich and Audiffren, 2011), which is based on a rebound surge of oxygenated blood post-exercise to the prefrontal cortex resulting in long-term destabilization of brain function. While the reasons and mechanisms for these physiological changes in these brain areas remain uncertain, they point out that frontal, sensorimotor, and striatal brain areas are regions of interest for studies mapping the neural correlates of exercise.

Previous studies have focused on brain network changes following exercise and have reported mixed results with both increased and decreased functional connectivity within specific ICA-derived brain networks soon after a bout of exercise (Schmitt et al., 2019) (Herold et al., 2020). Further experiments are thus clearly needed to replicate these effects after different types of exercise. Further, it is important to extend prior studies by characterizing the dynamics of brain networks over time as recent work points out that they can provide complementary information to ‘static’ functional connectivity measures (Alonso Martínez et al., 2020; Cabral et al., 2017; Caetano et al., 2022). Finally, the existing studies have focused on brain changes immediately after exercise, but have not tested, whether there are delayed changes in brain functional profile. Many of the exercise-related physiological changes occur during the recovery from exercise and it is important to study if this also applies to the brain.

To gain a better understanding of the prolonged effects of physical exercise on brain dynamics, the current study measured a 6-minute resting state fMRI in 24 young males at baseline and about one hour after a 60-minute aerobic exercise challenge (Saanijoki et al., 2018a). Leading Eigenvector Dynamic analysis (LEiDA) was used to identify recurrent phase-locking patterns (PL states) in fMRI signals and measure differences in their probability of occurrence and transition profiles before and after exercise. Independent component analysis (ICA) was used to identify static brain networks and measured potential differences in activity profiles. This study was explorative with no predefined hypotheses, but based on prior literature, we expected to find exercise-related changes in brain networks encompassing frontal and striatal brain areas.

## METHODS

The study was conducted in accordance with the Declaration of Helsinki at the Turku PET Centre, University of Turku and Turku University Hospital (Turku, Finland). The Ethics Committee of the Hospital District of Southwest Finland approved the study protocol. Twenty‐four men met the eligibility criteria, signed ethics-committee-approved informed consent forms, and were admitted into the study (Table 1). This study includes participants from a pre-registered study “Molecular and Functional Neurobiology of Physical Exercise; “EXEBRAIN” (NCT02615756) (http://www.clinicaltrials.gov).

**Table 1.** Descriptive statistics of the participants.

|  | Mean | Std. Deviation |
| --- | --- | --- |
| Age (years) | 26.833 | 5.297 |
| Height (cm) | 181.667 | 6.696 |
| Weight (kg) | 77.688 | 8.827 |
| BMI (kg / m<sup>2</sup>) | 23.483 | 1.640 |
| Activity (min/week) | 253.750 | 144.058 |
| VO<sub>2</sub><sub>max</sub> (l/min) | 3.628 | 0.684 |
| VO<sub>2</sub><sub>max</sub> (ml/kg/min) | 47.154 | 8.286 |
| Lactate pre-exercise | 1.033 | 0.347 |
| Lactate post-exercise | 1.333 | 0.491 |
| PANAS positive pre-exercise | 29.500 | 4.540 |
| PANAS positive post-exercise | 31.125 | 4.972 |
| PANAS negative pre-exercise | 12.417 | 2.603 |
| PANAS negative post-exercise | 12.083 | 2.603 |
| Beck Depression Inventory (BDI-II) | 3.458 | 3.753 |

The inclusion criteria for the study were; sex male, age 18–65 years, BMI ≤ 27 kg/m^2^, and good health. The exclusion criteria were current medication affecting the central nervous system, history of current neurological or psychiatric disease, and any chronic medical defect or injury, which hindered or interfered with everyday life. For all subjects, the exclusion criteria also included regular use of tobacco products or illicit drugs, heavy alcohol consumption, poor compliance, history of other nuclear imaging studies, and presence of ferromagnetic objects or significant claustrophobia, that would contraindicate MR imaging.

### The experimental setup

The experimental design has been previously described (Nummenmaa et al., 2018; Saanijoki et al., 2022, 2018a, 2018b). The participants were invited to positron emission tomography (PET) and MRI scans on two separate days (in a counterbalanced order). On one of the visits, we acquired baseline neuroimaging with no exercise, and on the other, the participants performed an exercise challenge before the PET scan that began within 15–36 minutes after the completion of the exercise session. After the PET scan the participants relocated to an MRI scanner, and the resting state fMRI scan used in the current study was acquired as the first sequence of the MRI session (70-90 minutes after exercise).

On the day of the baseline visit, the participants rested passively for 60 minutes before the scans without reading, music, television, or mobile entertaining devices. The exercise visit included 60 minutes of continuous aerobic cycling (Tunturi E85, Tunturi Fitness, Almere, The Netherlands) at a workload in the middle between aerobic and anaerobic thresholds predetermined individually in a maximal exercise test (described in detail in Saanijoki et al., 2018) (mean workload during exercise 157 (*SD* 47) W; 53 (*SD* 7)% from Load_max_; range 70– 265 W; average heart rate during exercise 144 (*SD* 15) beats per min; 74 (*SD* 7)% from maximal heart rate). Two participants had a slightly longer exercise session (76 and 77 min instead of programmed 60 min) due to an unexpected delay in the radiotracer supply. Music, television, or other technical devices were unavailable to the subjects during exercise. All participants completed the exercise session successfully.

### Questionnaire measurements

The Beck Depression Inventory (BDI-II) was used to measure current depressive symptoms. BDI-II is a 21-item self-reporting questionnaire for evaluating the severity of depression in normal and psychiatric populations (Beck et al., 1988; Jackson-Koku, 2016). Subjective feelings of pleasant versus unpleasant emotions were measured using the Positive and Negative Affect Schedule (PANAS) (Watson et al., 1988) before and after exercise.

### Physical activity and aerobic fitness measurements

Self-reported physical activity was assessed with a questionnaire in which participants rated the frequency (days/week) and duration (hours and minutes/week) of moderate-to-vigorous physical activity and other physical activity during the last three months. Fitness was evaluated as peak oxygen consumption (VO_2peak_), which was determined in a maximal exercise test performed on a cycle ergometer starting at 40-50 Watt (W) and followed by an increase of 30 W every 2 minutes until volitional exhaustion. Ventilation and gas exchange were measured (Jaeger Oxycon Pro; VIASYS Healthcare) and reported as the mean value per minute. The highest 1-min mean value of oxygen consumption was expressed as the VO_2peak_.

### MRI acquisition

Data were acquired with a 3-Tesla Philips Ingenuity PET-MR scanner at Turku PET Centre. Resting-state fMRI data were acquired with echo-planar imaging (EPI) sequence, sensitive to the BOLD signal contrast with the following parameters: TR=2000 ms, TE= 20 ms, 90° flip angle, 240 mm FOV, 80 × 80, 53.4 kHz bandwidth, 3 × 3 × 4 mm^3^ voxel size. Each volume consisted of 35 interleaved slices acquired in ascending order without gaps. A total of 180 functional volumes were acquired (total scan duration 6 minutes). Anatomical reference images were acquired using a T1-weighted sequence with 1 mm^3^ resolution (TR=8.1 ms, TE=3.7 ms, flip angle 7°, scan time ~5 minutes).

### fMRI preprocessing

Resting-state fMRI data were preprocessed using FSL 5.0.3 (FMRIB’s Software Library, www.fmrib.ox.ac.uk/fsl). Preprocessing steps included: the removal of the first five volumes of the acquisition to allow for signal stabilization/magnetization equilibrium; slice-timing correction using Fourier-space time-series phase-shifting; high-pass temporal filtering (Gaussian-weighted least-squares straight line fitting, with sigma=50.0s); motion correction with MCFLIRT (Jenkinson et al., 2002), including default rigid body alignment of every volume to the middle image of the acquisition and subsequent calculation of mean functional image and registration of individual images to the mean image; skull stripping of the functional data with the Brain Extraction Tool (BET) (Smith, 2002); spatial smoothing using a Gaussian kernel of FWHM 5mm; grand-mean intensity normalization of the entire 4D dataset by a single multiplicative factor. Resting-state fMRI scans were first registered to the respective T1 structural scan using boundary-based registration as implemented in FSL’s Linear Image Registration Tool (FLIRT) (Jenkinson et al., 2002) and then to a standard space template using non-linear registration (FNIRT) (Andersson et al., 2007).

After the basic preprocessing, FMRIB’s Independent Component Analysis (ICA)-based Xnoiseifier (ICA-FIX v1.068) was used to automatically denoise the data (Griffanti et al., 2014; Salimi-Khorshidi et al., 2014). Training data ‘Standard.RData’ was used with ICA classification threshold of 20 and high-pass filtering full width (2*50s) for motion confounds clean-up. Following the automated denoising, all registrations and FIX classifications were manually inspected. For FIX classifications, 70% of the data was checked randomly. No misclassified signal components were found during this inspection. The noise component classification was thus deemed successful for the current data.

### Dynamic Functional Connectivity with Leading Eigenvector Dynamics Analysis (LEiDA)

To measure dynamic functional connectivity, Leading Eigenvector Dynamics Analysis (LEiDA)(Cabral et al., 2017) approach was carried out in MATLAB R2018b (MathWorks, Natick, MA, USA). To apply LEiDA, average fMRI signals were obtained in 90 cortical and subcortical areas defined according to the Automated Anatomical Labelling (AAL) atlas using *fslmeants* from FSL. The time courses were then bandpass filtered (0.02 – 0.10 Hz); the analytic phase was obtained using the Hilbert transform, and the leading eigenvectors of the phase coherence matrices were calculated at each time point (Cabral et al., 2017). The eigenvectors were then clustered via K-means clustering using cosine similarity as distance, and optimizing across 200 replicates in line with prior work (Lord et al., 2019; Vohryzek et al., 2020).

For this explorative study, the number of clusters (K) was varied between 2 and 8, based on previous studies indicating that the optimal number of intrinsic networks is typically between 3 and 8 (Cabral et al., 2017, Lord et al., 2019, Vohryzek et al., 2020). Each K-means clustering yielded K clusters of patterns, where each cluster is represented by the average of all the patterns assigned to that cluster. These clusters represent distinct patterns in which brain areas are connected together in terms of phase alignment, and are referred to as phase-locking states or brain states (Hancock et al., 2022; Lord et al., 2019; Vohryzek et al., 2020). Two derived brain measures of interest were evaluated: probability of occurrence (the percentage of time points during which a certain brain state occurred) and switching probability (the likelihood of transitioning from a given state into any of the other states) (Vohryzek et al., 2020; (Marschall et al., 2023).

### Static Brain Connectivity with Independent Component Analysis (ICA)

Group ICA across both the baseline and the exercise fMRI data was performed using FSL’s MELODIC (Probabilistic independent component analysis for functional magnetic resonance imaging) with the number of components set to 25 (Schmitt et al., 2019). This was followed by dual regression that uses the group-level ICA maps to perform multivariate temporal regression of the individual component time courses to yield subject-specific spatial maps. Differences between these spatial maps was then investigated in a voxel-wise general linear model (GLM) that tests associations within each network in activation patterns across the measurement.

### Statistical analysis

### Baseline vs. post-exercise conditions

The descriptive statistics and correlations were estimated with JASP (Version 0.16.4). The main statistical model compared baseline and post-exercise conditions within subjects. LEiDA models for probability of occurrence and between network transitions were tested with a two-tailed paired t-test using 5000 permutations tests in Matlab. Multiple comparisons correction was performed with FDR q < 0.05 and Bonferroni corrections. The group ICA brain networks were used in dual regression that includes statistical inference with a general linear model (GLM; ‘paired t-test contrast’) and FSL randomize for which multiple comparison corrections were performed with 500 permutations and threshold-free cluster enhancement (TFCE) corrected p < 0.05 (Smith and Nichols, 2009).

### Exploratory correlations analyses

Given the exploratory nature of the study, we were utilized correlation analysis to map associations between demographics, questionnaire data (Table 1) and brain measures that show statistical between condition differences without formal correction for multiple comparisons.

## RESULTS

### Exercise destabilizes striato-fronto-parietal brain state

We found that the probability of occurrence of a striato-fronto-parietal network was significantly reduced in the post-exercise condition compared to baseline, with a concomitant increase in the probability of occurrence of a pattern of global functional connectivity, where no functional network is activated (Figure 1, Table 2). This striato-fronto-parietal brain state is consistently detected when varying the number of clusters K between 2 and 8, showing statistically significant differences in occupancy between baseline and exercise condition (p < 0.05), but only survives multiple comparisons correction (FDR q < 0.05) when considering up to 4 clusters/states. (Supplement 1). Brain areas in the implicated brain state included the bilateral caudate, putamen, and pallidum, as well as the middle orbitofrontal, frontal operculum, frontal trigonum, and inferior parietal cortices (Figure 1, Supplement 2).

**Table 2.** Descriptive statistics for probabilities of occurrence. P values are based on paired sample t-test and permutation testing over 5000 permutations.

| Descriptive Statistics |  |  | Baseline vs. post exercise |
| --- | --- | --- | --- |
|  | Mean | Std. Deviation | p |
| State #1 baseline | 0.608 | 0.165 | p = 0.0135 |
| State #1 post exercise | 0.687 | 0.140 |  |
| State #2 baseline | 0.156 | 0.085 | p = 0.0170 |
| State #2 post exercise | 0.126 | 0.079 |  |
| State #3 baseline | 0.142 | 0.073 | p = 0.1359 |
| State #3 post exercise | 0.105 | 0.066 |  |

**Figure 1.**
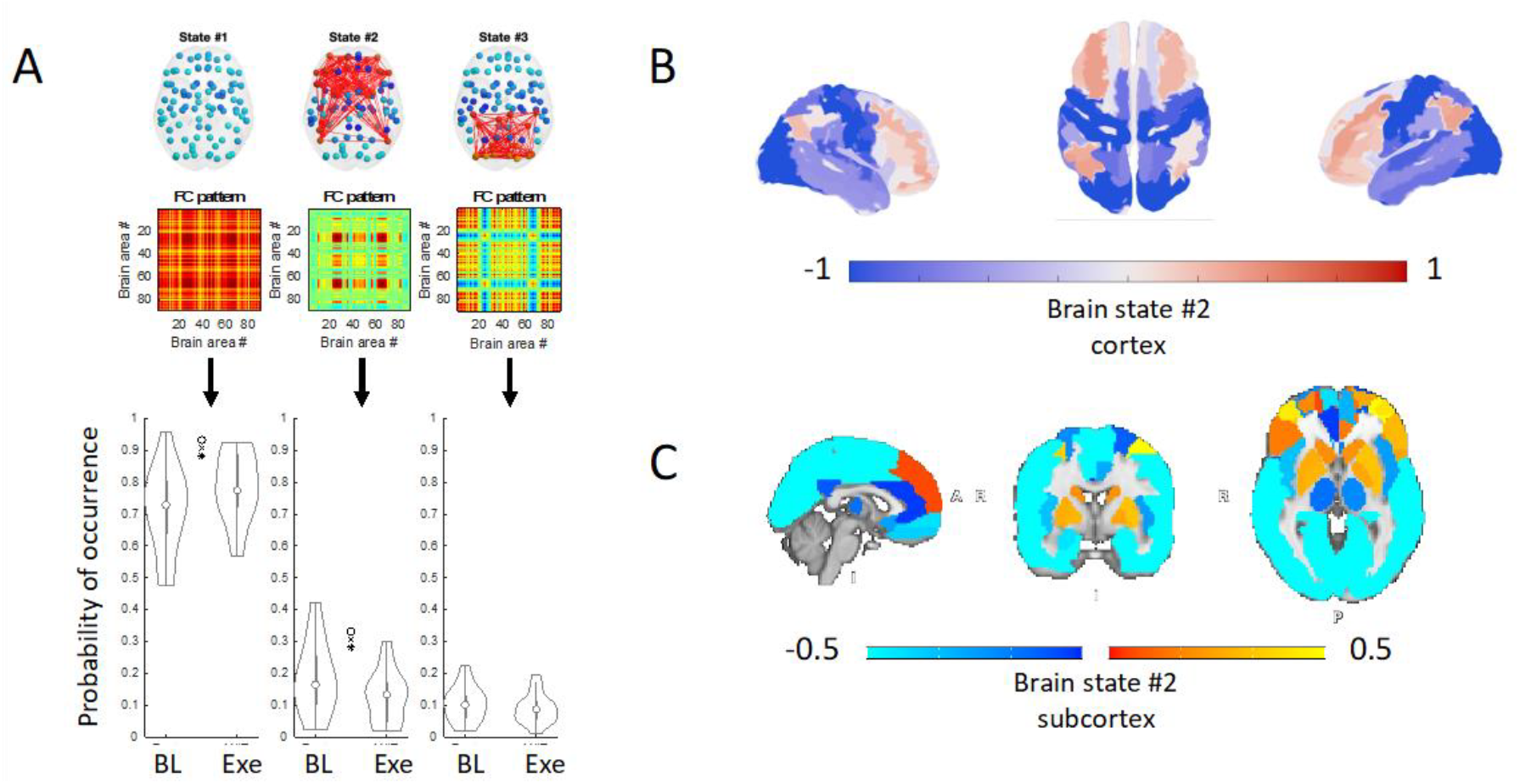
The destabilization of brain network dynamics in striatal and frontoparietal brain areas when comparing baseline (BL) to exercise condition ca. 1h after cycling exercise challenge as compared to the baseline measurements. A) The cluster centroids representing the three brain states are shown on a glass brain where? the network of brain areas are synchronizing together (top plot), as well as the corresponding phase coherence matrices representing the pattern of functional connectivity (FC) between brain areas (middle plot), as well as violin plots of the distribution of state probabilities of occurrence (bottom plot). The probabilities of occurrence were statistically compared between pre and post exercise using a permutation-based paired t-tests over 5000 permutations, with (*) indicating uncorrected p-value < 0.05; (x) FDR-corrected q < 0.05; and (o) Bonferroni corrected p < 0.05). B) Visualization of the brain areas in the network of interest brain state #2 showing the cortical distribution of the network. C) Visualization of the brain areas in brain state #2 showing the subcortical distribution of the network. Note that we use different colors and plotting range in B and C to improve distinction between brain areas.

### Exercise-related differences in brain states have minor influence on network transitions

The brain state to brain state switching profiles were different between the baseline and post-exercise conditions, but the differences were not statistically significant (Figure 2). This additional analysis implicates that exercise increased the probability of occurrence of the ‘global brain state’ and decreased the probability of occurrence of the striato-fronto-parietal brain state without affecting the overall dynamic patterns of brain activity.

**Figure 2.**
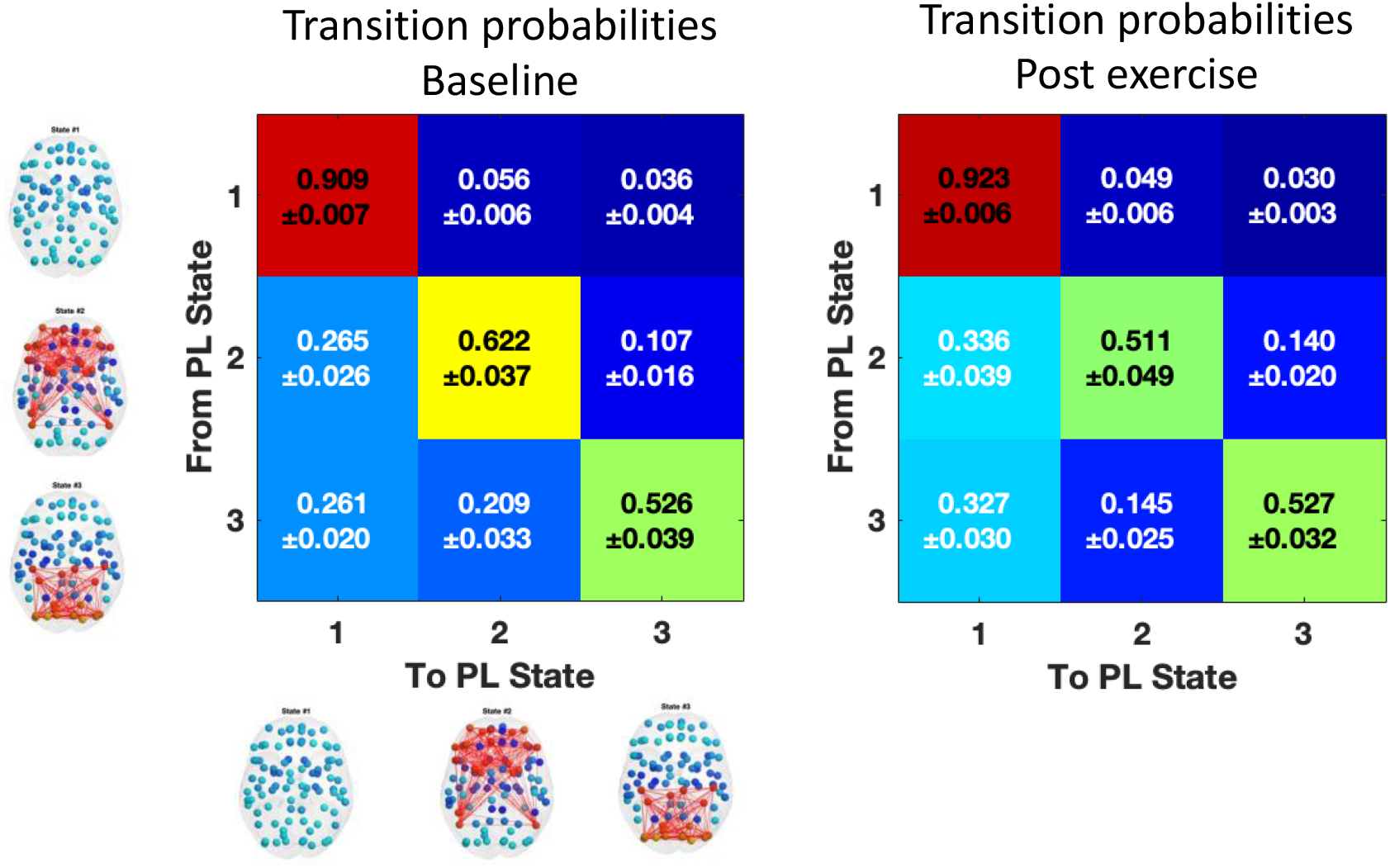
The transition probabilities between different brain states. The matrices show the mean probabilities and standard deviations for remaining in a brain state (diagonal) and transitioning to any of the other brain states both at baseline condition (left) and exercise condition (right). The transition profiles showed minor variance between conditions, but the differences were not statistically significant.

### ICA-derived brain networks are not modulated by exercise

We detected canonical resting-state networks across the group ICA across fMRI data from the baseline and exercise conditions across all participants (Eyre et al., 2021); Rajasilta et al., 2020), and 11 networks were identified (Figure 3) and compared between conditions using dual regression. No statistically significant differences between baseline and exercise conditions were found.

**Figure 3.**
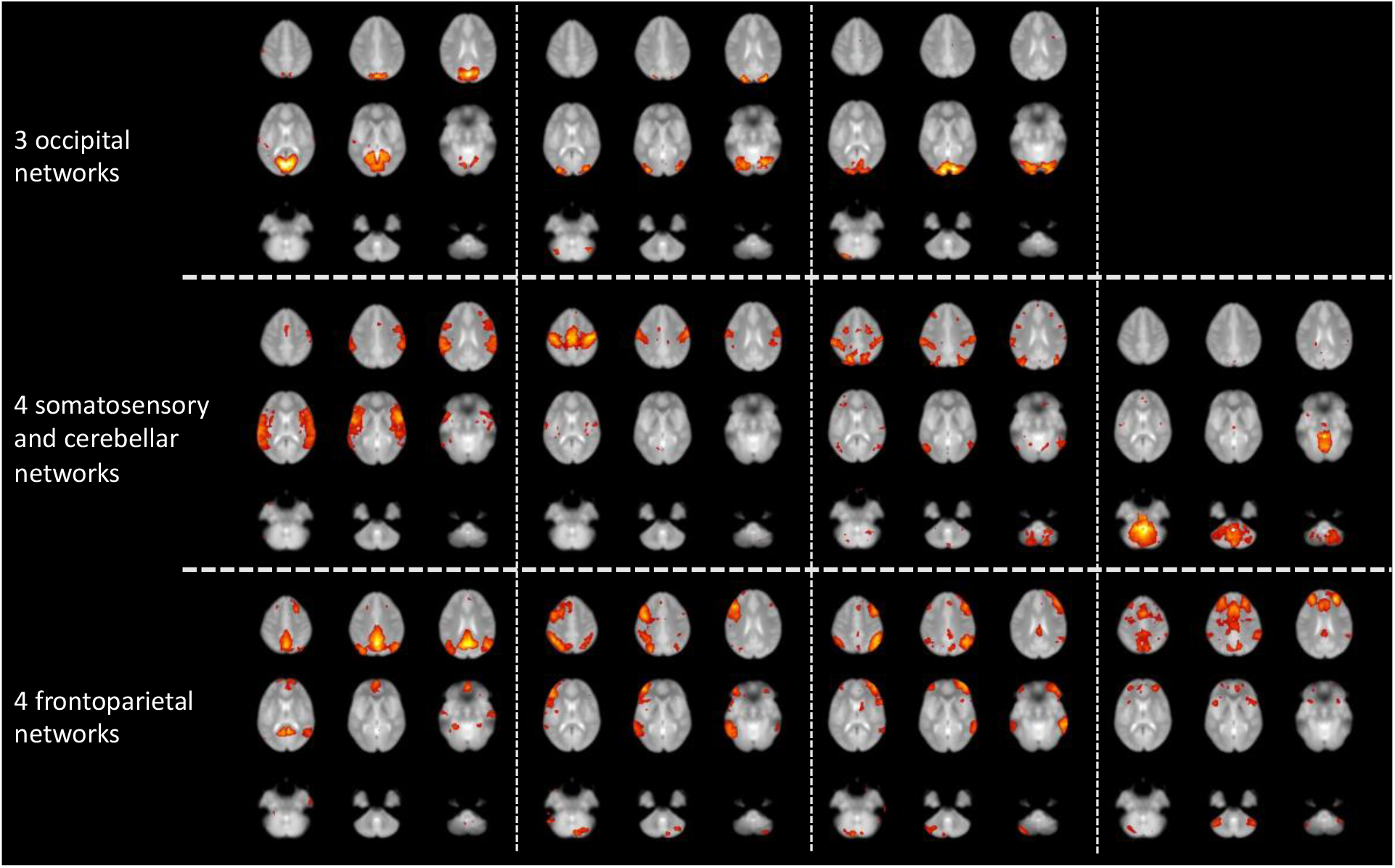
The 11 brain networks identified in ICA analyses. Each network is represented in 9 axial slices. None of the identified networks showed statistically significant differences between baseline and post-exercise conditions.

### Exploratory correlation analyses

Correlations of brain network probabilities of occurrence and demographics are provided in Supplementary Tables 1 and 2. The correlations were not statistically significant except for pre-exercise lactate levels and the probability of occurrence of brain state #1 in post-exercise scans (Spearman’s rho = 0.468, p = 0.021) (Supplementary Tables 1 and 2).

## DISCUSSION

Our main new finding was that the post-exercise state was associated with reduced periods of synchronization between striatal, frontal, and parietal regions and increased periods of global synchronization. Given that increased global synchronization has been previously found to be significantly related with higher cognitive task performance in healthy older adults (Cabral et al., 2017) and to be reduced in patients with schizophrenia (Farinha et al., 2022), and that reduced connectivity between the frontal and subcortical brain areas has been linked with lower scores of perceived stress (Caetano et al., 2022). Together these findings suggest that the exercise-induced network changes observed here may be directly associated with known positive effects of physical exercise in brain function. Moreover, the fact that no statistically significant differences were detected following exercise using ICA analysis that does not take into account the temporal evolution of functional connectivity, reinforce the need to include the temporal dimension in the analysis of resting-state fMRI signals.

There is mounting evidence that exercise training in older adults improves functional connectivity (FC) within several resting-state networks, that is altered during both healthy and pathological aging (Burdette et al., 2010; Voss et al., 2010). In addition, a single acute bout of exercise in young adults can lead to increased connectivity in sensorimotor resting networks (Rajab et al., 2014). However, these prior studies assumed that the functional connectivity between distinct regions of interest remains constant in a task-free setting. While this assumption offers a useful framework for analysis and interpretation, it fails to account for the dynamic and adaptive nature of human brain activity. Furthermore, considerable within-subject variation in FC has been observed within a single imaging session where changes in FC strength and direction were noted in timescales of minutes or even seconds. In the current study, we used the relative phase of the fMRI signals (as in LEiDA), where the modeling is sensitive to the underlying network patterns with the advantage that the networks can have spatial overlap and occur transiently at different frames in time (Marschall et al., 2023). Other available approaches for dynamic connectome modeling include the sliding window technique, time-frequency analysis, dynamic connectivity regression, and dynamic connectivity detection (Hutchison et al., 2013). Despite the interesting findings yielded by studies using dynamic modeling (Kaiser et al., 2016; Pan et al., 2022; Zhao et al., 2022), some controversy exists regarding the methodology (Hindriks et al., 2016), confounding effects (Nikolaou et al., 2016), and reliability (Zhang et al., 2018) of dynamic functional connectivity.

Significant changes in brain hemodynamics in response to acute physical exercise have been observed and some of these can be attributed to increases in blood flow during and immediately after exercise (Smith and Ainslie, 2017). In addition, changes in circulating metabolites, such as lactate, may improve the brain energy metabolism and function (Tsukamoto et al., 2016). While these factors may directly affect brain activity, near-infrared spectroscopy (NIRS) and fMRI studies have established robust post-exercise changes in brain activity, which can be discerned from physiological changes alone (Herold et al., 2018; Yu et al., 2021). Still, given that the destabilization of the striato-fronto-parietal network was observed at the expense of increased stability of a state of global synchrony, it is possible that the impact of exercise in brain function is directly linked with global changes in physiological processes interacting with the brain.

Since resting fMRI data in the current study were acquired after the cessation of cycling exercise and the subsequent PET scanning (approx. 60 mins), physiological changes pertaining to perfusion and cardiac output unlikely explain the results. However, comparisons to studies that focused on acute and prolonged physical exercise and brain function are not straightforward due to influence of different factors such as exercise protocols (Mehren et al., 2019), cardiorespiratory fitness level (Li et al., 2019), sex of the participants (Mehren et al., 2019), and time delays between the cessation of exercise and cognitive testing (Chang et al., 2012).

A recent meta-analysis of 20 studies investigated the influence of physical exercise on cognition-related functional brain activation (Yu et al., 2021). The findings suggested that physical exercise interventions lead to changes in functional activation patterns primarily located in the precuneus and those associated with frontoparietal, dorsal attention, and default mode networks. The precuneus plays a key role in the frontoparietal network by interconnecting parietal and prefrontal regions and thus has a strong engagement in core cognitive domains, including attention and executive functions (Bullmore and Sporns, 2009). In addition, dynamic network models may enable novel means to explore the hypofrontality hypothesis (Loprinzi et al., 2019). The hypofrontality hypothesis specifically entails that prefrontal cortex-dependent tasks (e.g., working memory tasks) may be compromised during high-intensity exercise. While the current study did not include cognitive measurements, the brain areas that jointly showed a change in dynamics have been implicated in cognitive processing, and future studies could test whether brain dynamics mediate acute and longer-term links between exercise and cognition.

The study has several limitations. The current study aimed to capture the effects of exercise at the recovery phase after about 60 minutes. The inclusion of more repeated measurements is needed in the future studies. Age and sex of the participants can influence biophysical determinants of acute exercise. The current study only included healthy males with good fitness levels, whereas a more diverse group, including females, more variable fitness levels, and age groups should be considered. The study only employed a rather short fMRI sequence with limited time points (180 volumes) to control the overall scan session time, and future studies may benefit from longer and/or faster fMRI sequences.

In conclusion, dynamic modeling was used to ascertain how exercise affects the brain’s repertoire of functional network states under task-free conditions. The study showed that dynamic brain states are sensitive and could capture theoretically relevant changes in striatal and frontoparietal networks in the recovery phase of a 60-minute bout of exercise, which provides a unique viewpoint on the literature by pointing out neural changes in the recovery phase post exercise. This provides new insights for future studies with repeated neuroimaging and coupled cognitive measures to unravel the network-level neural underpinnings of exercise.

## Acknowledgements

We thank all participants that took part in the studies, and Turku PET center staff for carrying out the measurements together with the investigators.

## Disclosure of interest

The authors report no conflict of interest.

## Data availability statement

The Finnish law and ethical permissions do not allow the sharing of the data used in this study.

## Author contributions

- **Jetro J. Tuulari**, participated to the health screening of the participants, conceptualized the resting state fMRI data analyses, performed data preprocessing and modeling, performed the statistical analyses, wrote the initial draft of the manuscript.
- **Wajiha Bano**, participated to critical evaluation of the used preprocessing, performed statistical analyses and wrote the initial draft of the manuscript.
- **Tiina Saanijoki**, collected the behavioral and neuroimaging data.
- **Joana Cabral**, provided key expertise in creating the code and optimal settings for the LEiDA modelling.
- **Henrique Fernandes**, provided key expertise in fMRI data preprocessing and ICA models.
- **Marina Charquero Ballester**, provided key expertise in LEiDA modelling.
- **Jakub Vohryzek**, provided key expertise in LEiDA modelling.
- **Lauri Tuominen**, planned the study, participated to the health screening of the participants, implemented the used MRI sequences.
- **Lauri Nummenmaa**, planned the study, implemented the used MRI sequences.
- **Kari Kalliokoski**, conceived the study, and obtained funding.
- **Morten L. Kringelbach**, provided supervision on data analyses and the infrastructure for the work carried out during the fellowship of JJT.
- **Jussi Hirvonen** conceived the study, obtained funding, and provided supervision.
- **All authors** reviewed and approved the manuscript.

## Funding

- Paavo Nurmen Säätiö;
- Opetus-ja KulNuuriministeriö;
- Novo Nordisk Foundation;
- Suomen Kulttuurirahasto;
- Veritas Foundation;
- Paulon Säätiö;
- Instrumentariumin Tiedesäätiö;
- Sigrid Juséliuksen Säätiö;
- University of Turku Doctoral Programme of Clinical Research;
- Juho Vainion Säätiö;
- Medical Imaging Centre of Southwest Finland;
- Suomen Akatemia, Grant/Award Numbers: 251125, 251399, 256470, 265917, 281440, 283319, 304385;
- Turku Collegium for Science Technology and Medicine
- Suomen Lääketieteen Säätiö
- Emil Aaltosen säätiö
- Hospital District of Southwest Finland State Research Grants
- Signe and Ane Gyllenbergin Säätiö

## Notes

### Competing Interest Statement

The authors have declared no competing interest.

