## Supplementary figures and images for "Physical exercise and brain network dynamics: reduction of frontoparietal-striatal connectivity following 1 hour of aerobic cycling"

### Supplement 2

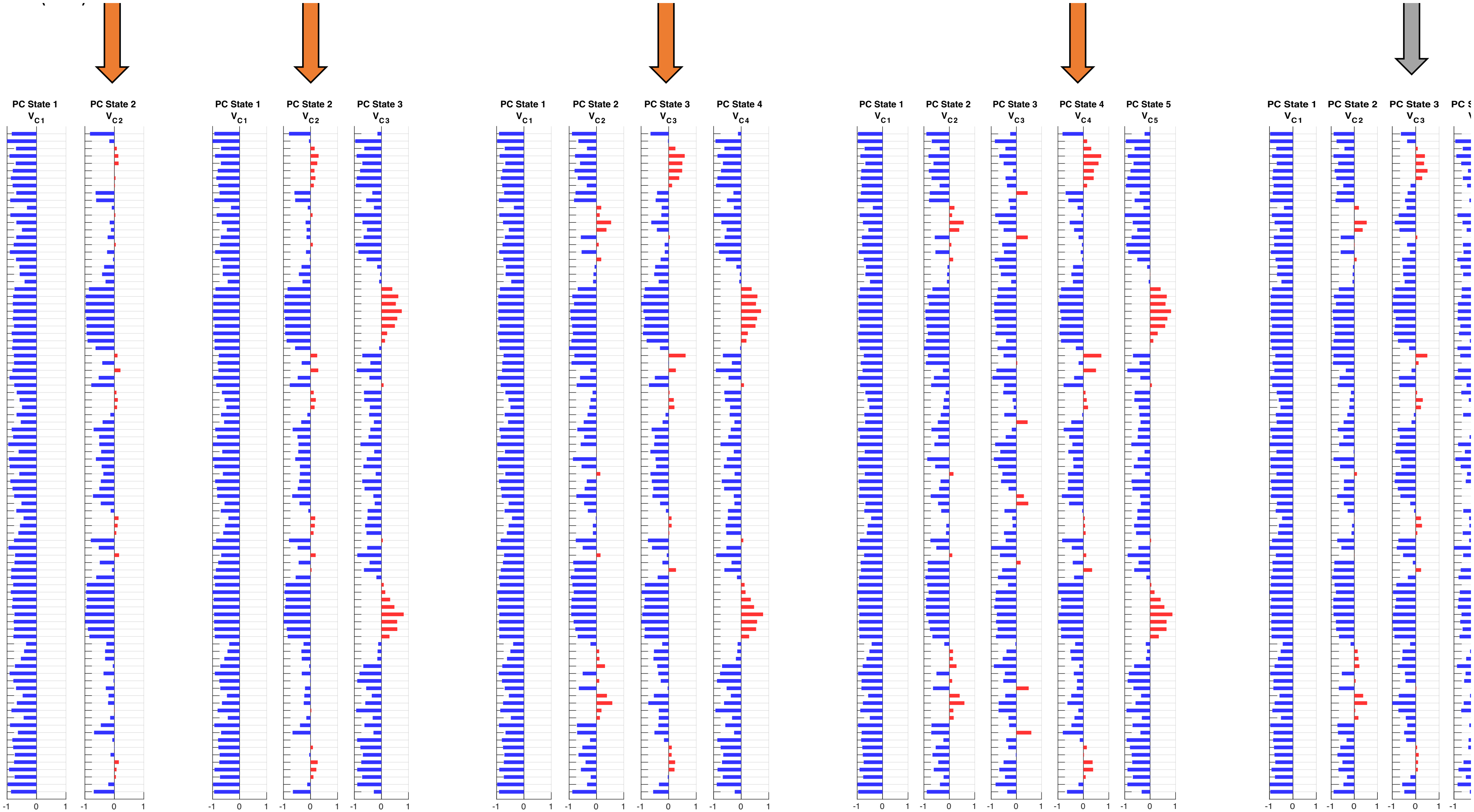
