## Supplement 1 for "Physical exercise and brain network dynamics: reduction of frontoparietal-striatal connectivity following 1 hour of aerobic cycling"

- Clustering solutions from K2 to K8 are shown with network visualization on the class brain, phase coherence matrices and violin plots of the distribution of state probabilities of occurrence.

- Permutation based paired t-test over 5000 permutations statistics are as follows:

  * = raw p-value < 0.05,

  x = FDR q < 0.05,

  o = Bonferroni corrected p < 0.05.

- The fronto-striato-parietal network of interest is marked with an orange asterisk *

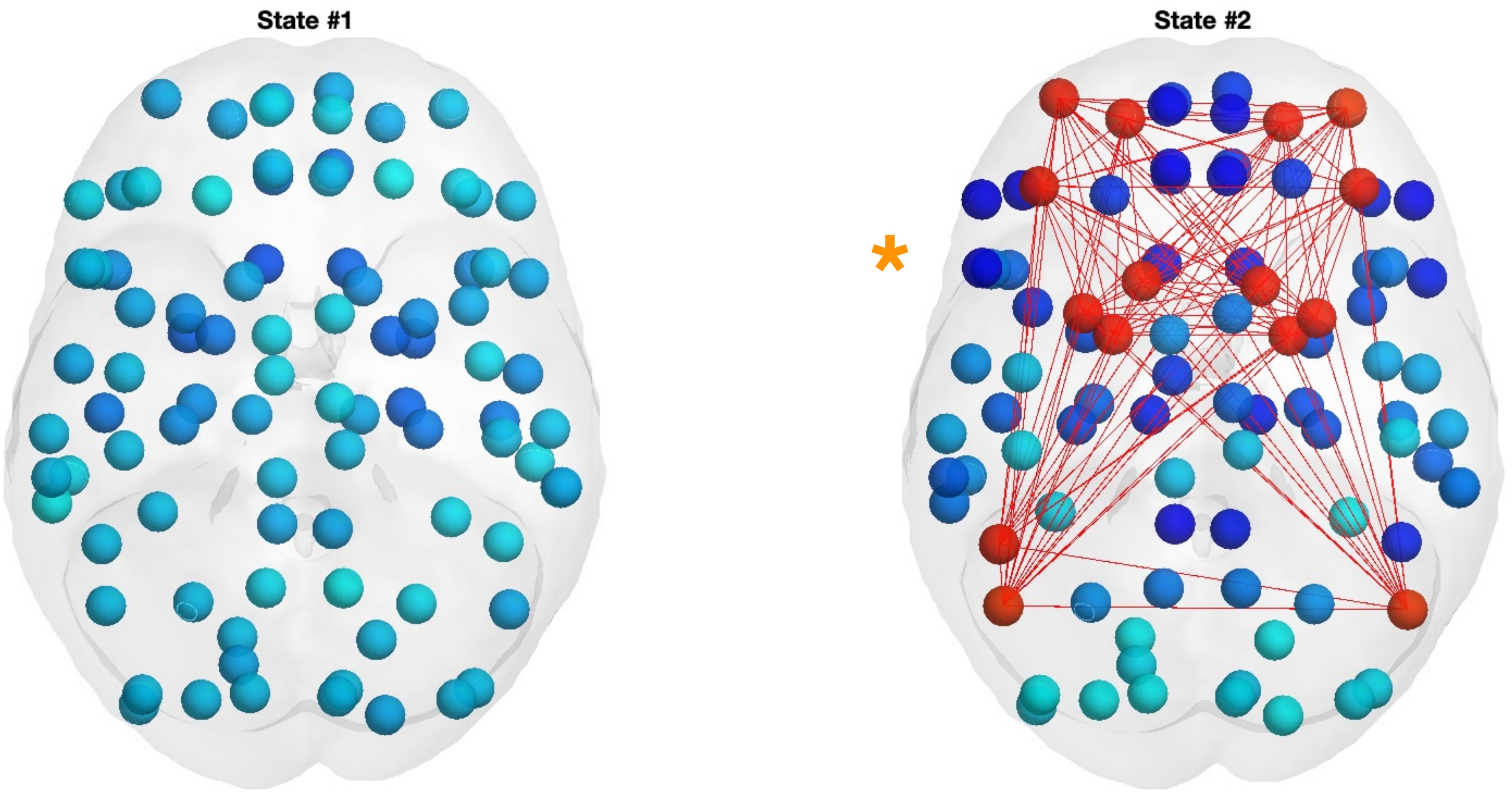
State #1
State #2
*

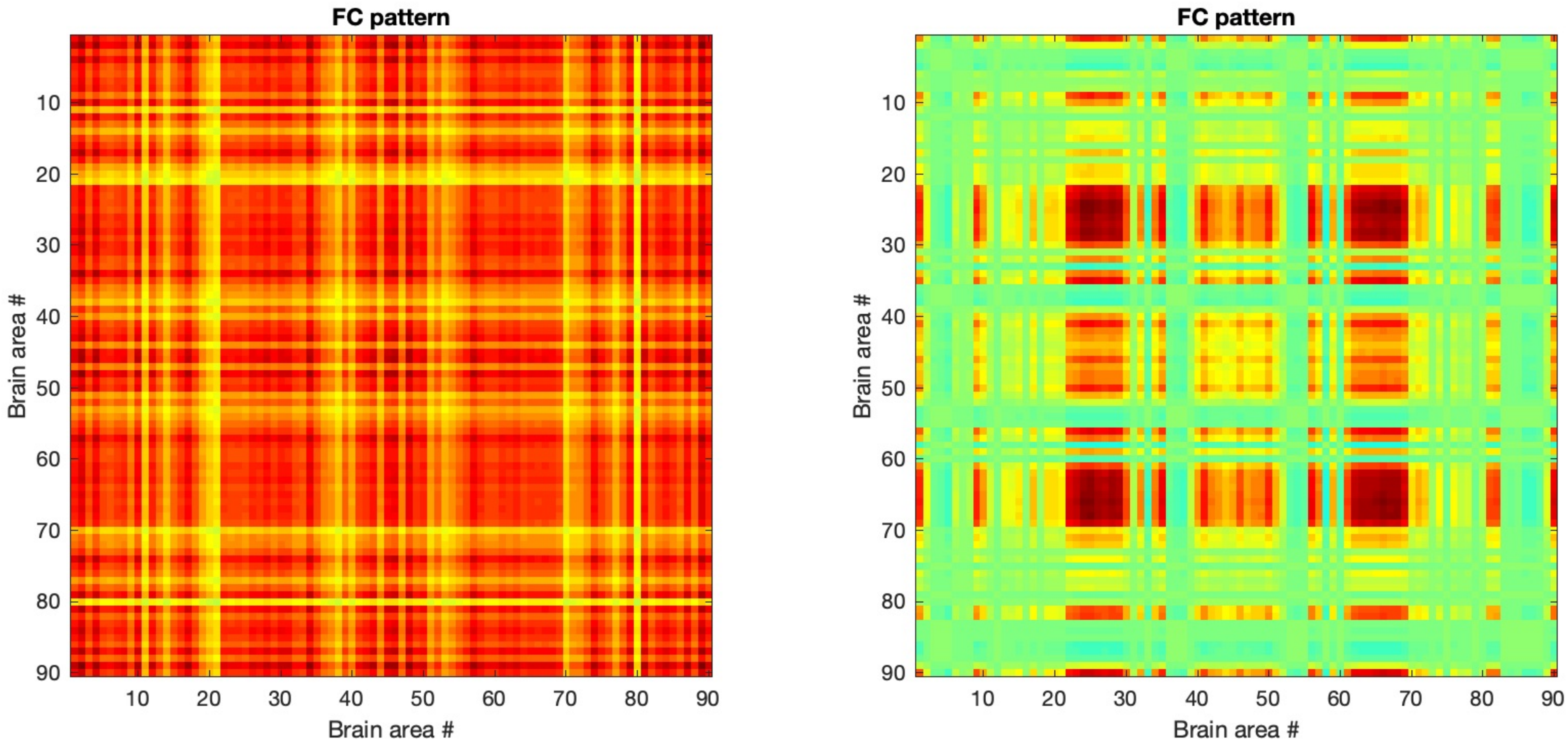
FC pattern
Brain area #
10
20
30
40
50
60
70
80
90
FC pattern
Brain area #

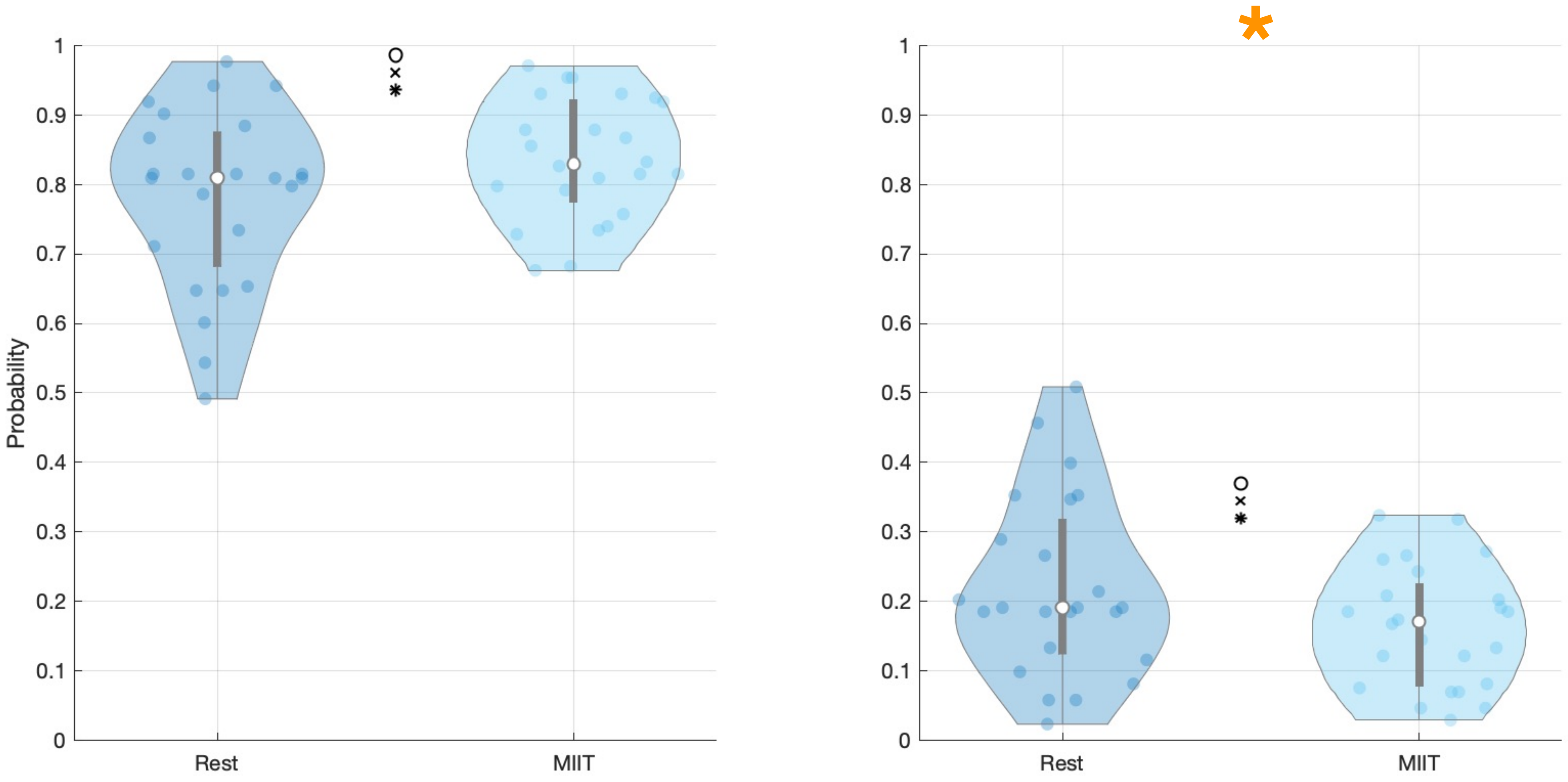

Probability
1
0.9
0.8
0.7
0.6
0.5
0.4
0.3
0.2
0.1
0
Rest
MIIT
*

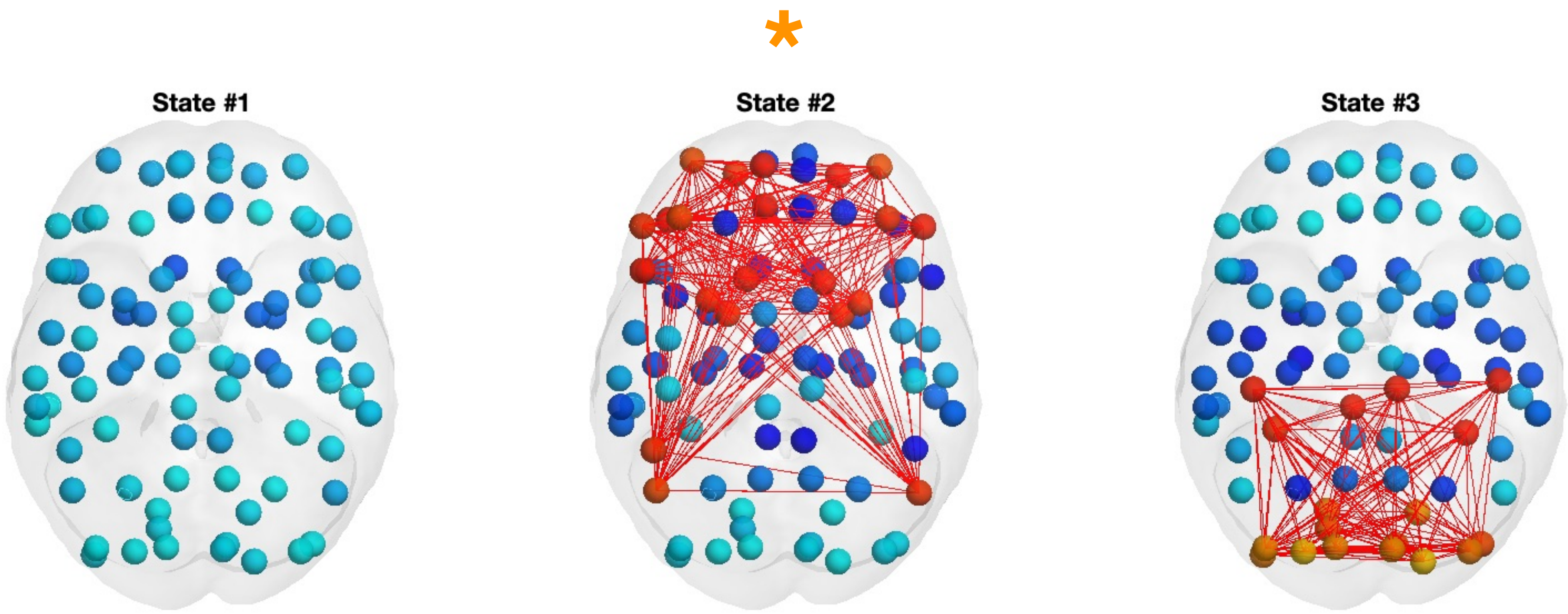
*
State #1
State #2
State #3

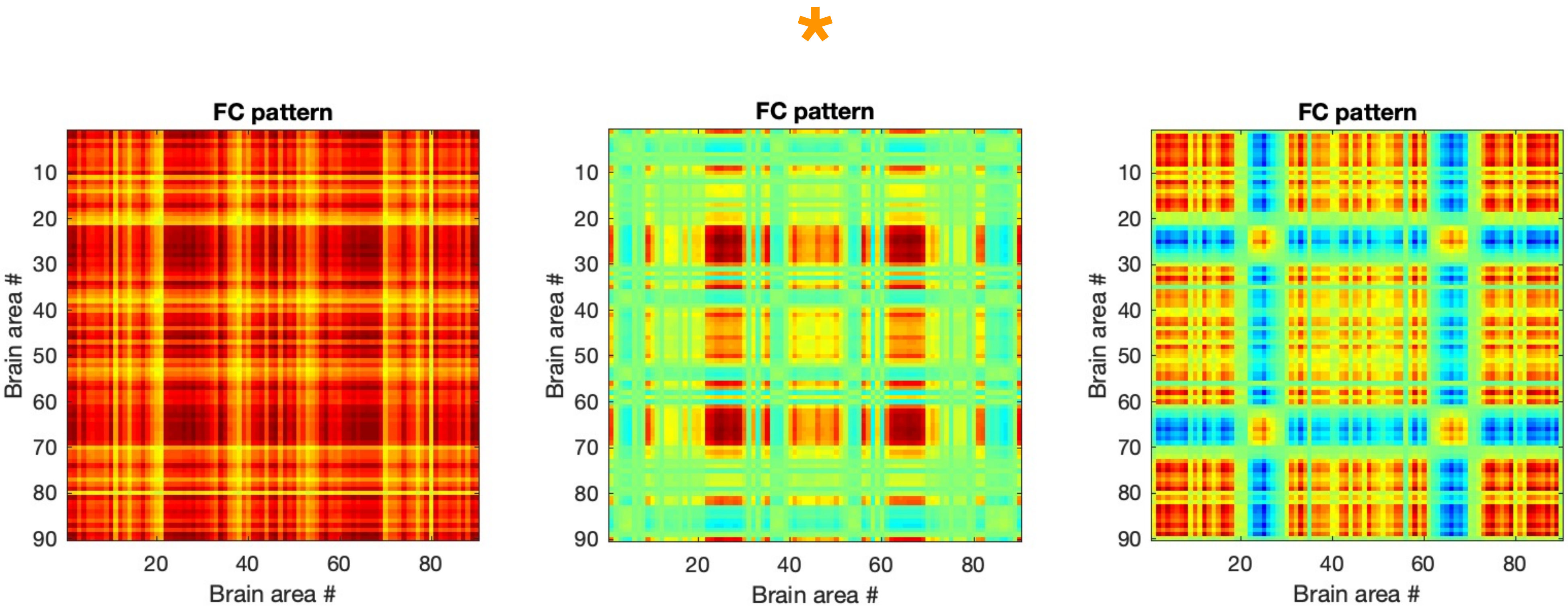
FC pattern
Brain area #
10
20
30
40
50
60
70
80
90
20
40
60
80
Brain area #
FC pattern
Brain area #
10
20
30
40
50
60
70
80
90
20
40
60
80
Brain area #
FC pattern
Brain area #
10
20
30
40
50
60
70
80
90
20
40
60
80
Brain area #

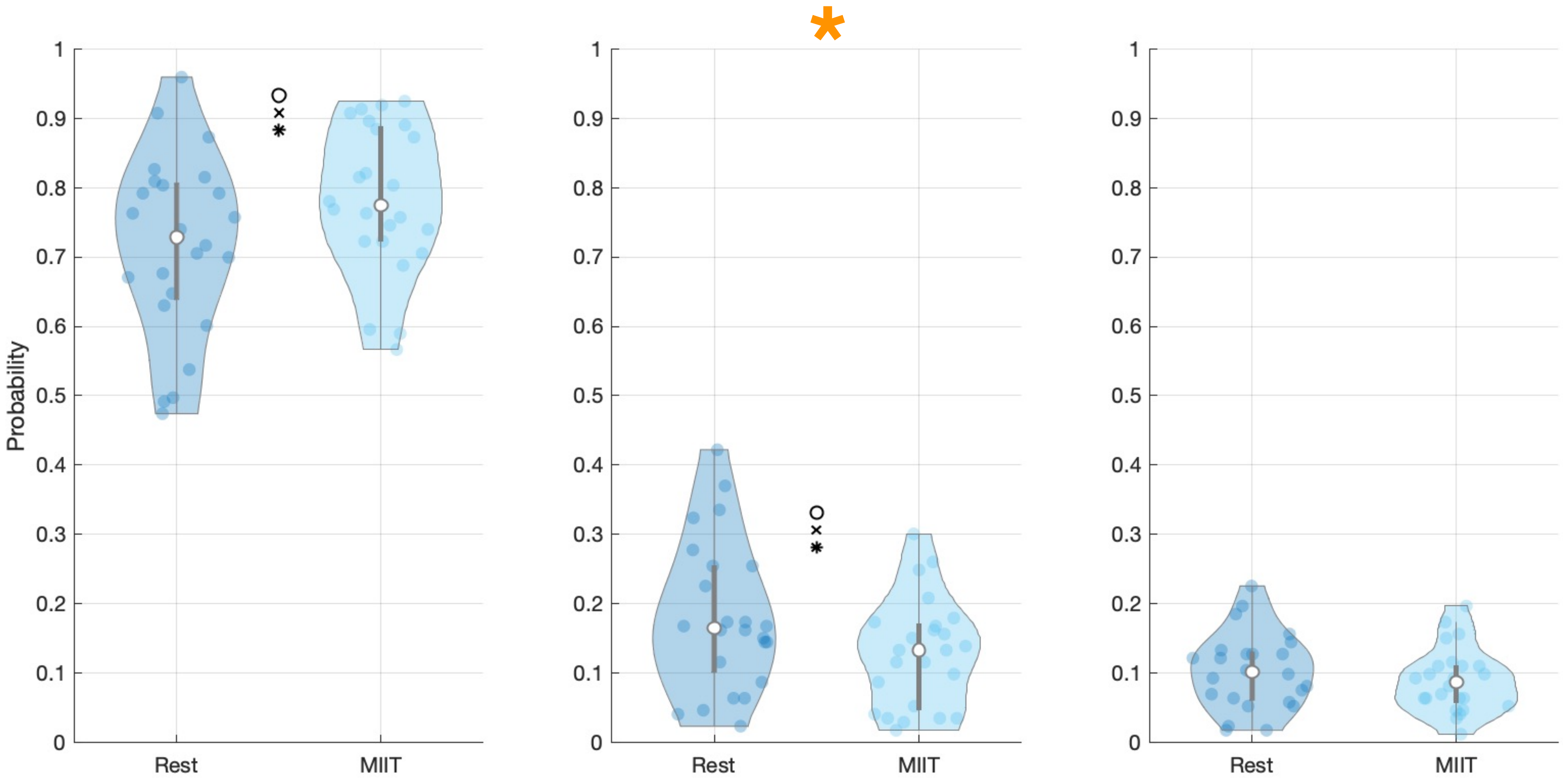

Probability
1
0.9
0.8
0.7
0.6
0.5
0.4
0.3
0.2
0.1
0
Rest
MIIT
*

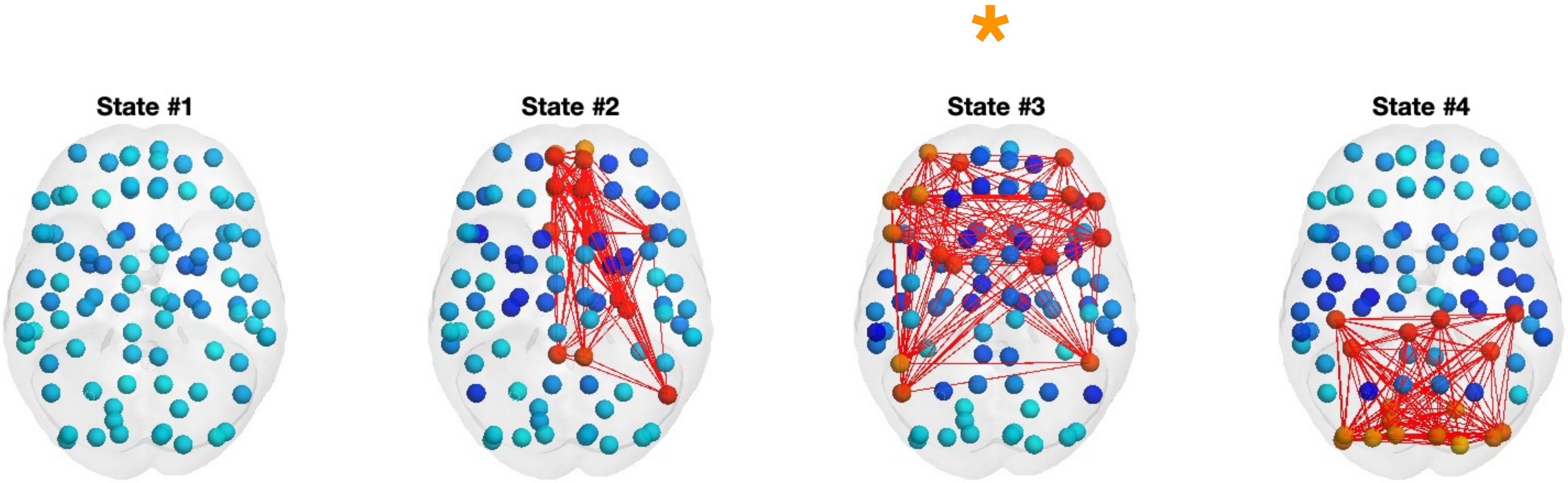
*
State #1
State #2
State #3
State #4

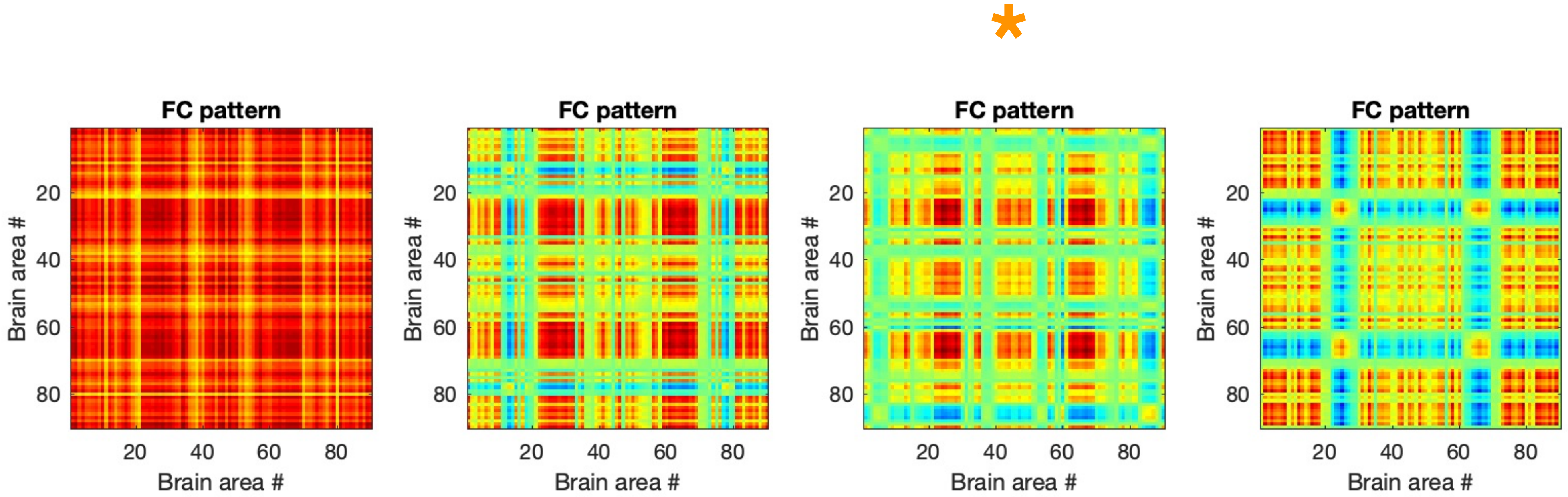
*
FC pattern
FC pattern
FC pattern
FC pattern
Brain area #
20
40
60
80

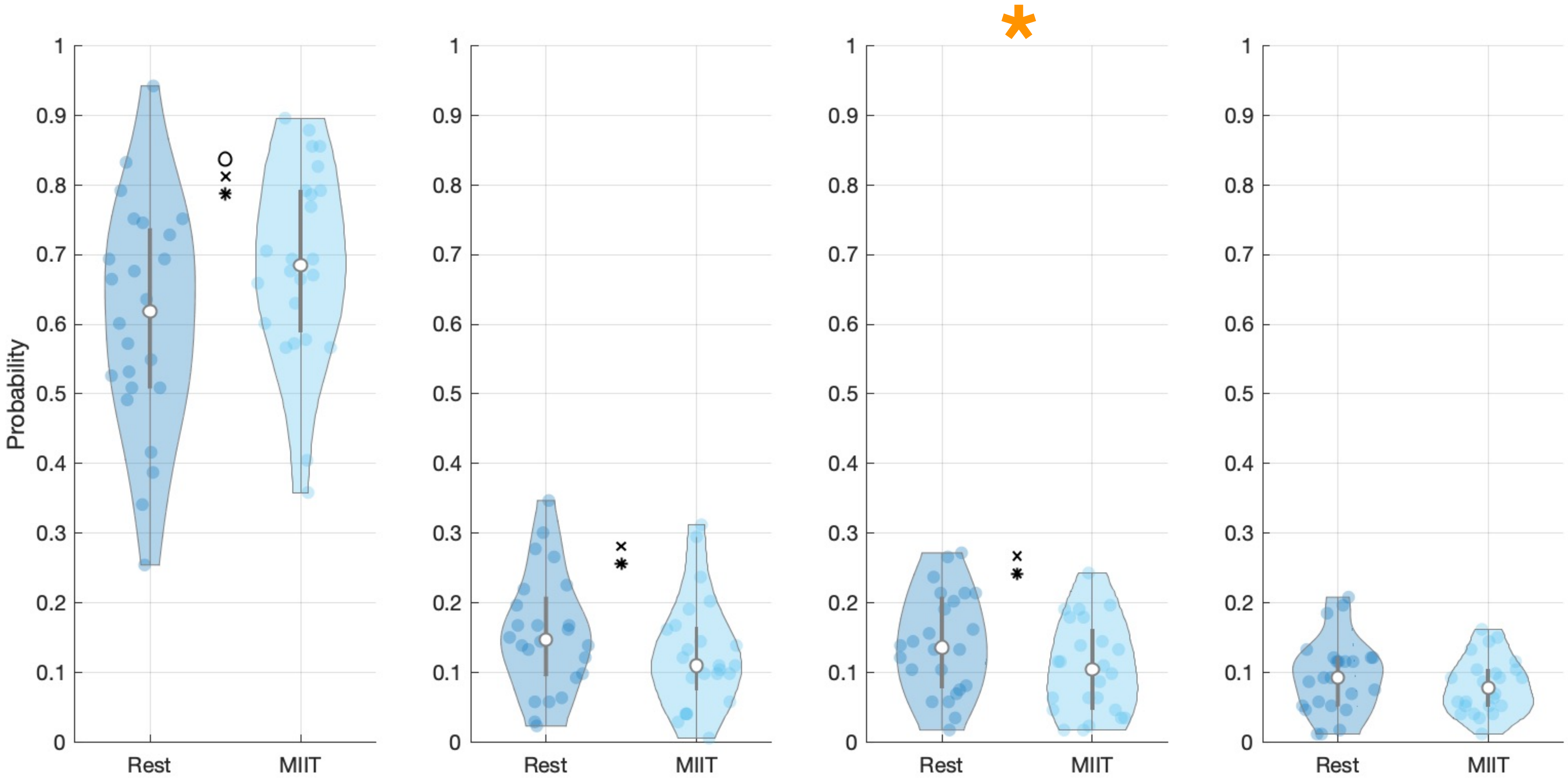

Probability
1
0.9
0.8
0.7
0.6
0.5
0.4
0.3
0.2
0.1
0
Rest
MIIT

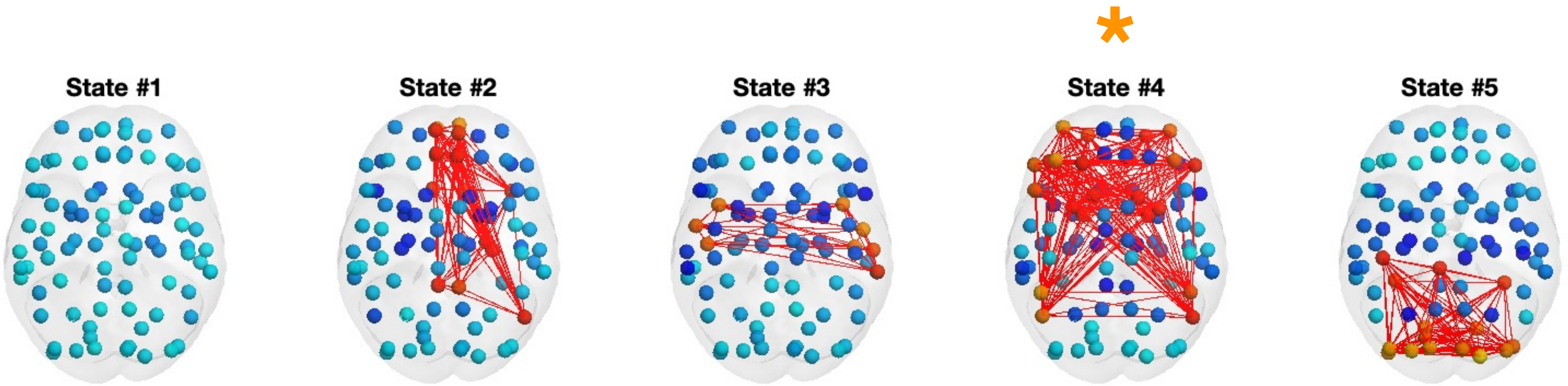

*
State #1
State #2
State #3
State #4
State #5

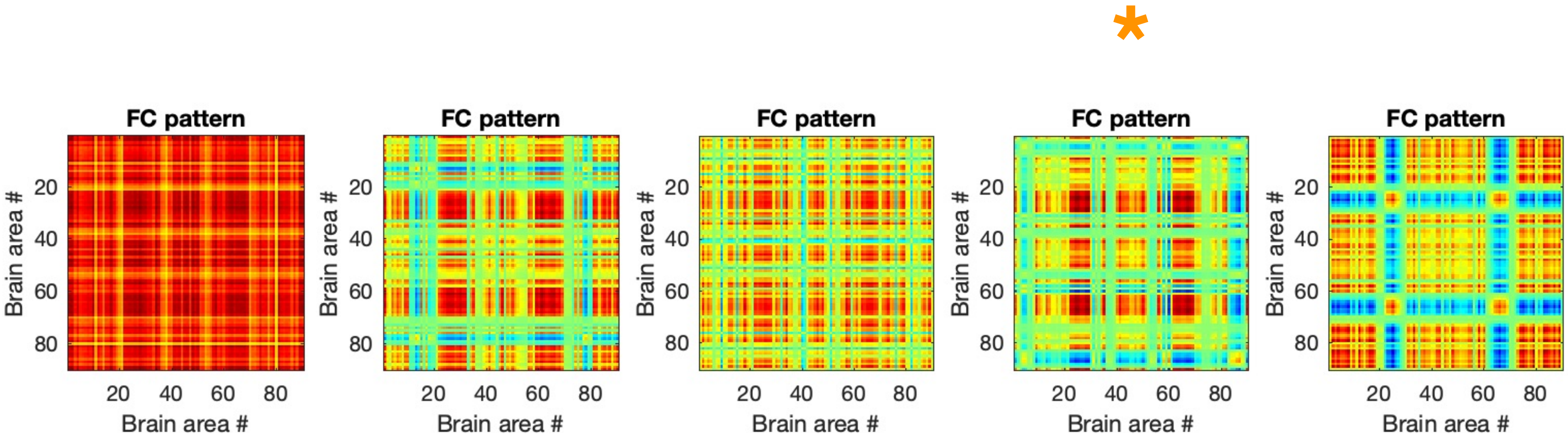
*
FC pattern
Brain area #
20
40
60
80

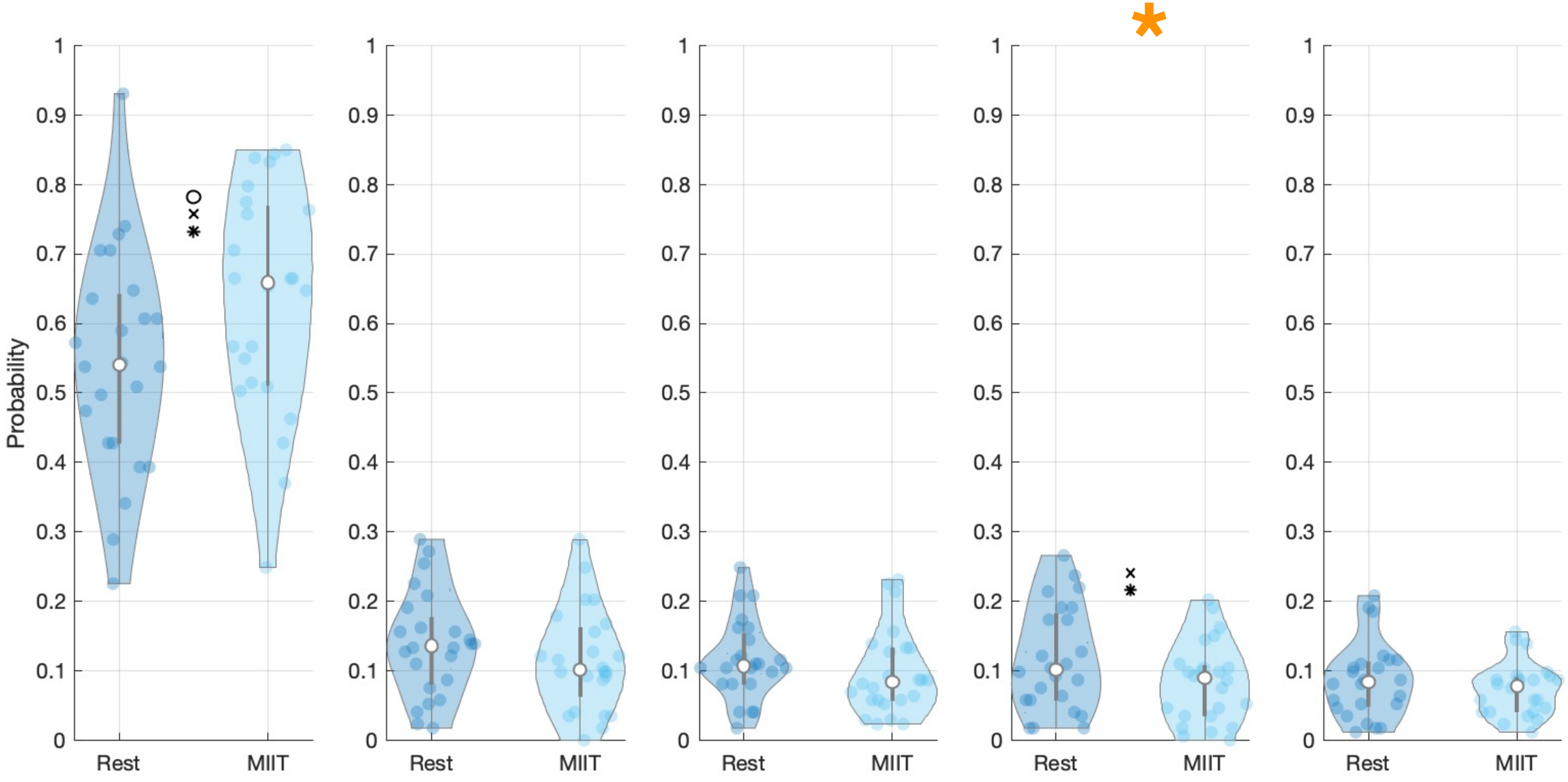

Probability
1
0.9
0.8
0.7
0.6
0.5
0.4
0.3
0.2
0.1
0
Rest
MIIT
*

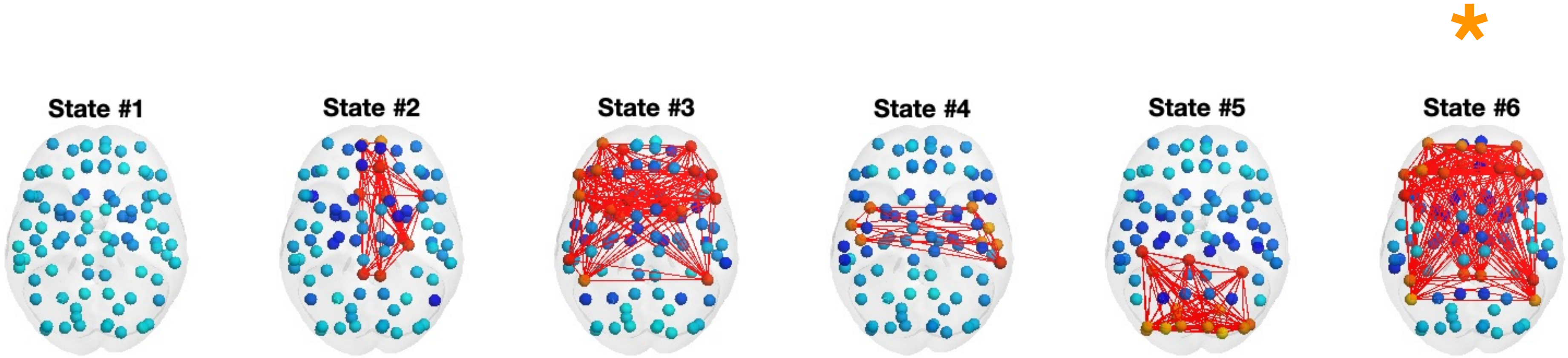
*
State #1
State #2
State #3
State #4
State #5
State #6

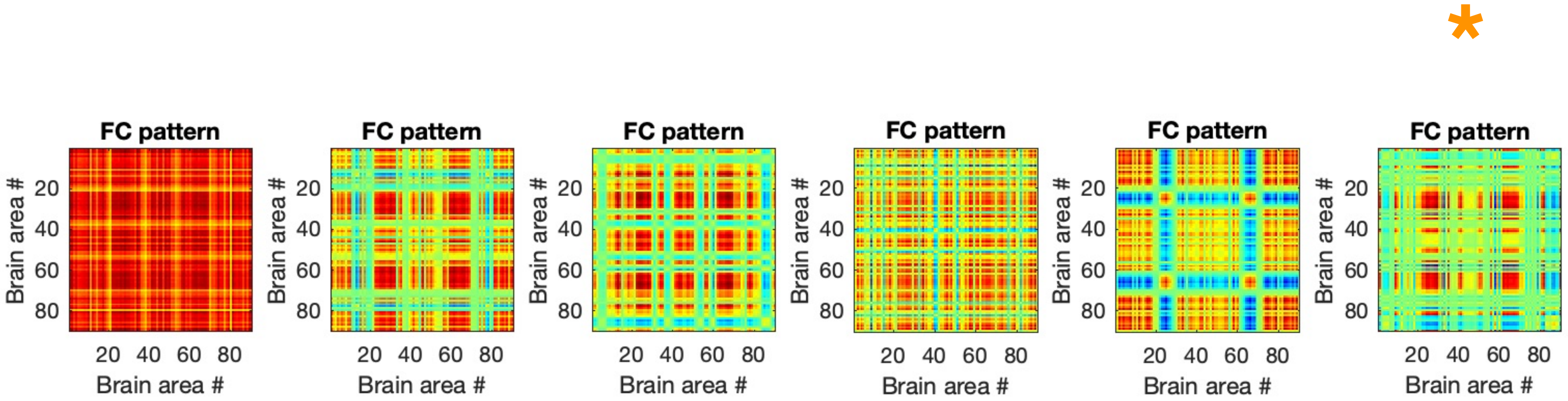
*
FC pattern
FC pattern
FC pattern
FC pattern
FC pattern
FC pattern
Brain area #
20
40
60
80
Brain area #

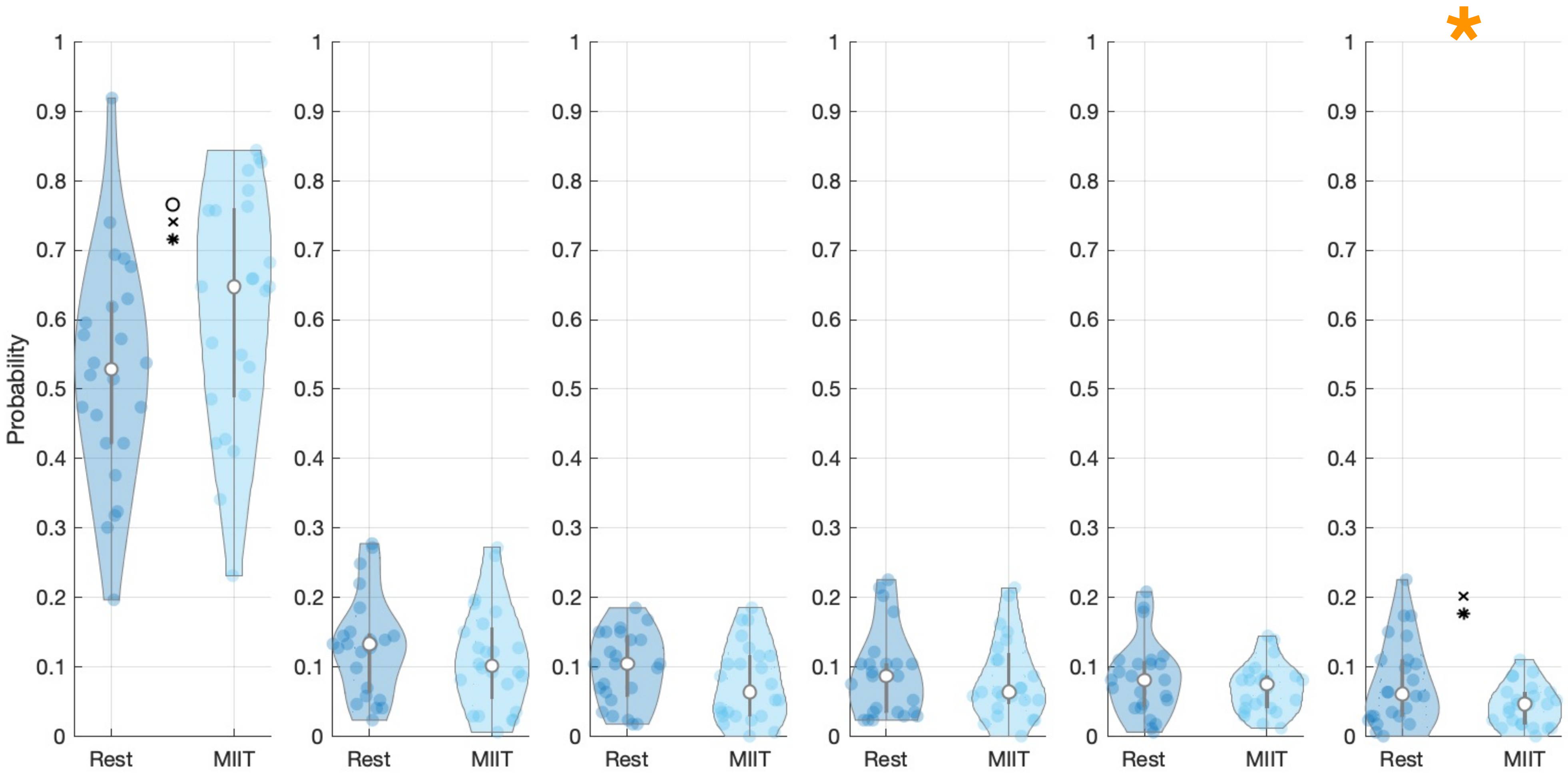

Probability
0
0.1
0.2
0.3
0.4
0.5
0.6
0.7
0.8
0.9
1
Rest
MIIT

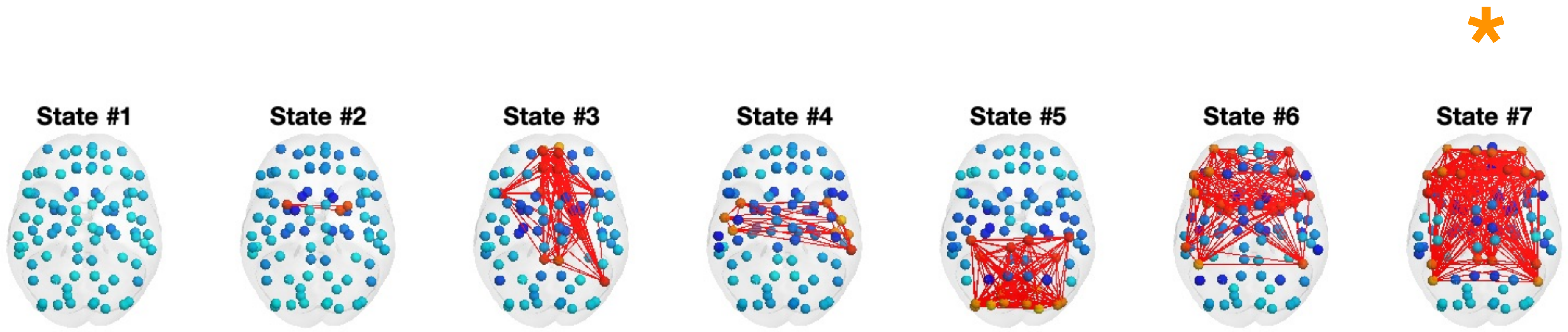
*
State #1
State #2
State #3
State #4
State #5
State #6
State #7

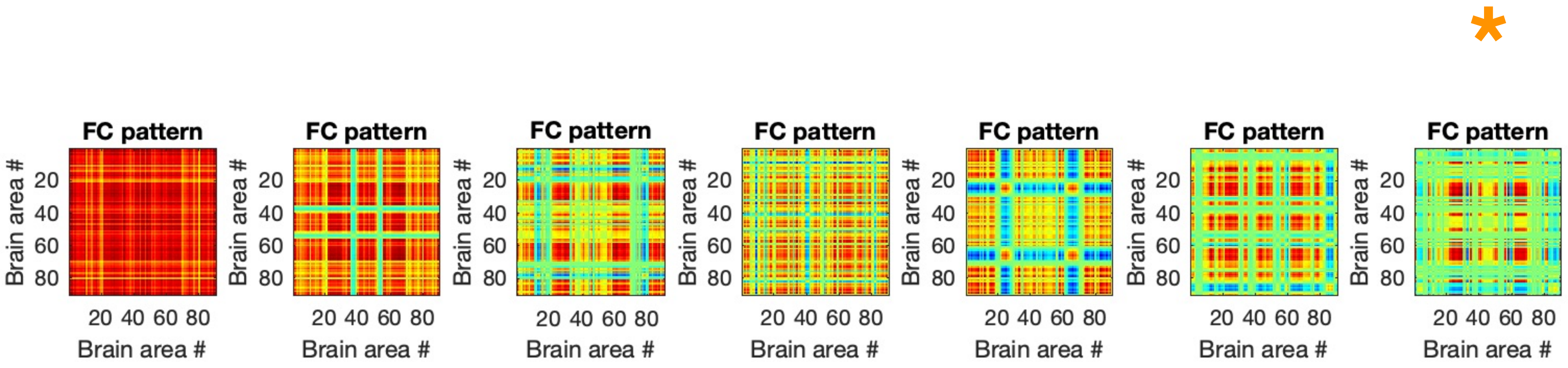

*
FC pattern
FC pattern
FC pattern
FC pattern
FC pattern
FC pattern
FC pattern
Brain area #
20
40
60
80
20 40 60 80
Brain area #

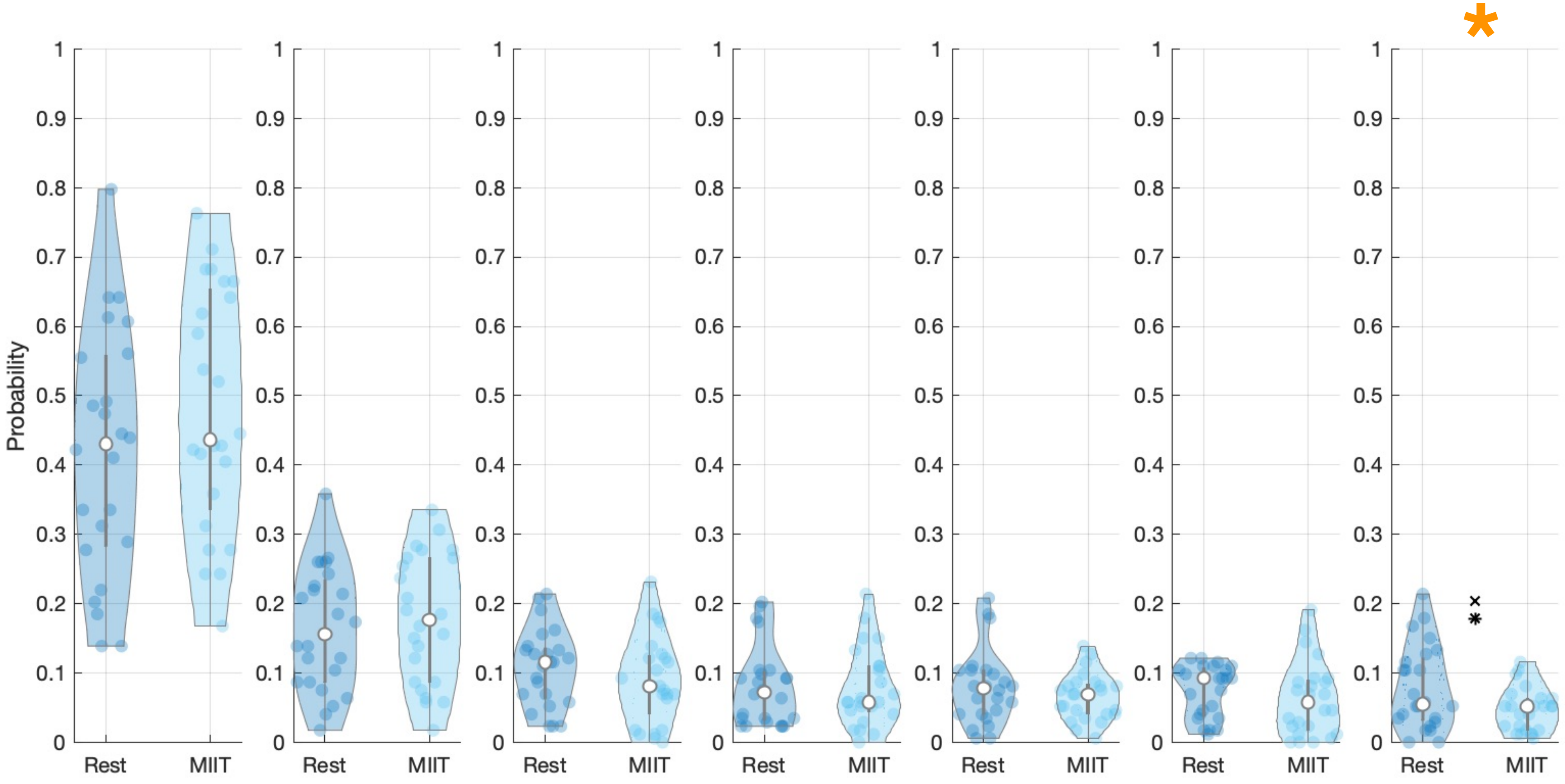

Probability
1
0.9
0.8
0.7
0.6
0.5
0.4
0.3
0.2
0.1
0
Rest
MIIT
*

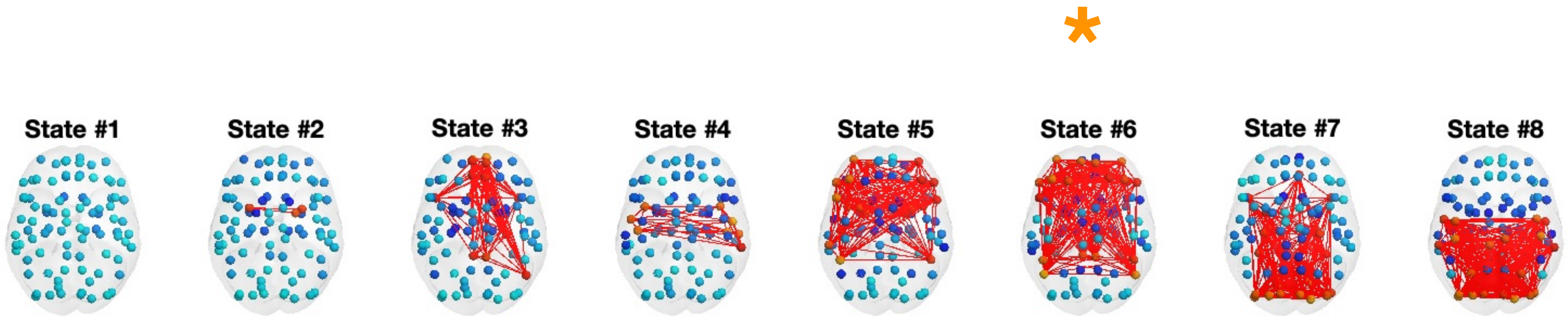
*
State #1
State #2
State #3
State #4
State #5
State #6
State #7
State #8

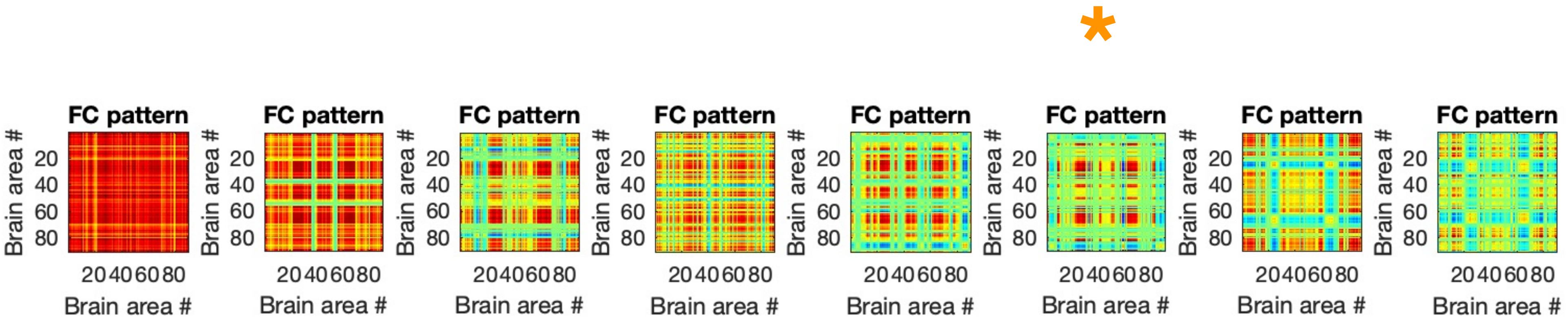
*
FC pattern
Brain area #
20
40
60
80
20 40 60 80
Brain area #
FC pattern
Brain area #
20
40
60
80
20 40 60 80
Brain area #
FC pattern
Brain area #
20
40
60
80
20 40 60 80
Brain area #
FC pattern
Brain area #
20
40
60
80
20 40 60 80
Brain area #
FC pattern
Brain area #
20
40
60
80
20 40 60 80
Brain area #
FC pattern
Brain area #
20
40
60
80
20 40 60 80
Brain area #
FC pattern
Brain area #
20
40
60
80
20 40 60 80
Brain area #
FC pattern
Brain area #
20
40
60
80
20 40 60 80
Brain area #

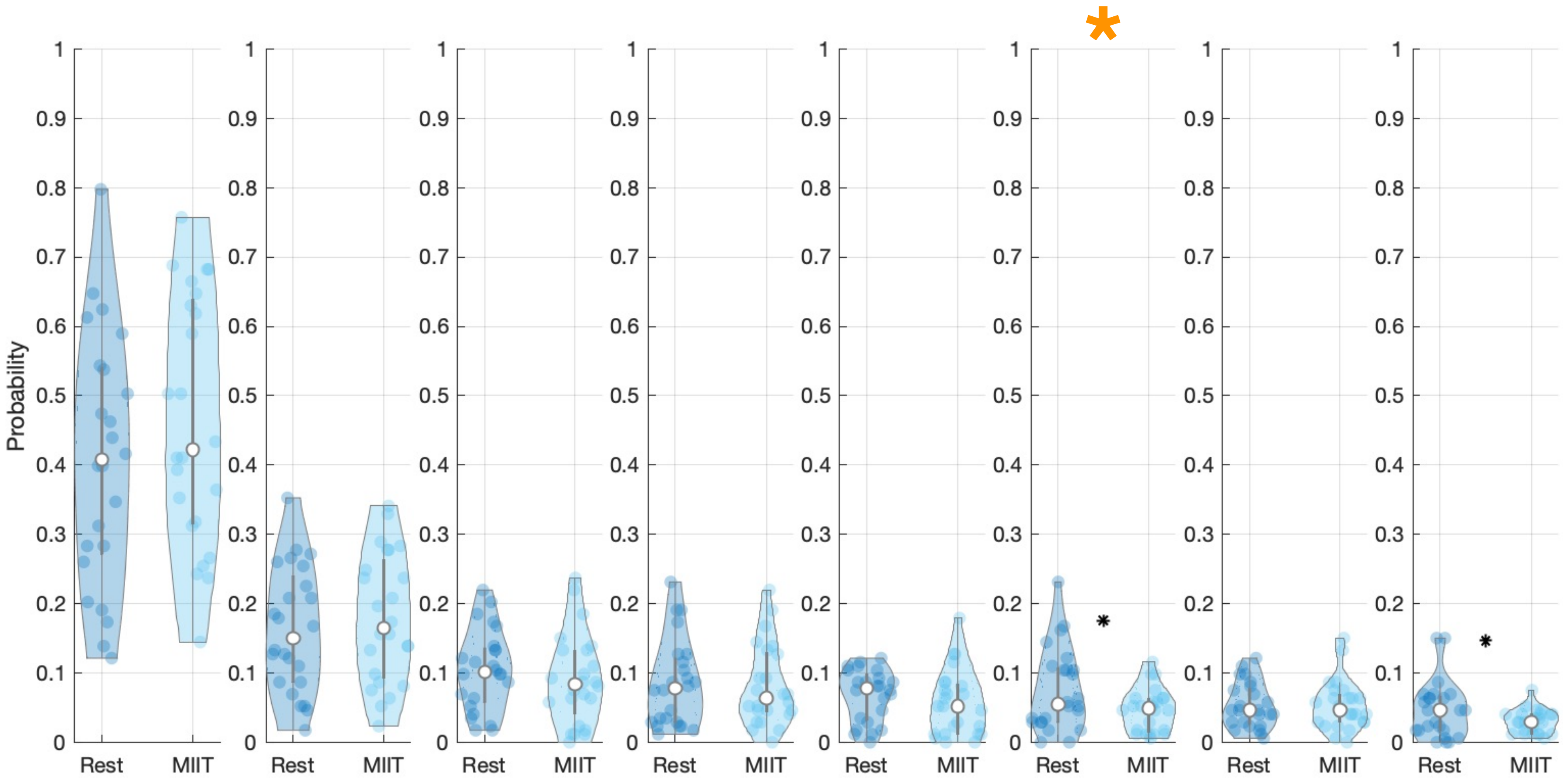

Probability
1
0.9
0.8
0.7
0.6
0.5
0.4
0.3
0.2
0.1
0
Rest
MIIT
*
