## Supplementary Table 1 for "Physical exercise and brain network dynamics: reduction of frontoparietal-striatal connectivity following 1 hour of aerobic cycling"

**Correlations baseline**

| Spearman's Correlations | | | | | | | | | | | | | | | | |
| --- | --- | --- | --- | --- | --- | --- | --- | --- | --- | --- | --- | --- | --- | --- | --- | --- |
| **Variable** | |  | | **1** | | **2** | | **3** | | **4** | | **5** | | **6** | | **7** |
| 1. Probability of occurrence brain state #1 baseline |  | Spearman's rho |  | — |  |  |  |  |  |  |  |  |  |  |  |  |
|  |  | p-value |  | — |  |  |  |  |  |  |  |  |  |  |  |  |
| 2. Probability of occurrence brain state #2 baseline |  | Spearman's rho |  | -0.836 | *** | — |  |  |  |  |  |  |  |  |  |  |
|  |  | p-value |  | < .001 |  | — |  |  |  |  |  |  |  |  |  |  |
| 3. Probability of occurrence brain state #3 baseline |  | Spearman's rho |  | -0.880 | *** | 0.711 | *** | — |  |  |  |  |  |  |  |  |
|  |  | p-value |  | < .001 |  | < .001 |  | — |  |  |  |  |  |  |  |  |
| 4. BMI |  | Spearman's rho |  | 0.050 |  | 0.117 |  | 0.084 |  | — |  |  |  |  |  |  |
|  |  | p-value |  | 0.816 |  | 0.585 |  | 0.696 |  | — |  |  |  |  |  |  |
| 5. Acitivity (min/week) |  | Spearman's rho |  | 0.092 |  | -0.178 |  | -0.062 |  | -0.232 |  | — |  |  |  |  |
|  |  | p-value |  | 0.669 |  | 0.406 |  | 0.773 |  | 0.274 |  | — |  |  |  |  |
| 6. VO2max (l/min) |  | Spearman's rho |  | -0.209 |  | 0.294 |  | 0.264 |  | 0.096 |  | 0.357 |  | — |  |  |
|  |  | p-value |  | 0.327 |  | 0.163 |  | 0.213 |  | 0.655 |  | 0.087 |  | — |  |  |
| 7. VO2max (ml/kg/min) |  | Spearman's rho |  | -0.213 |  | 0.155 |  | 0.085 |  | -0.539 | ** | 0.530 | ** | 0.667 | *** | — |
|  |  | p-value |  | 0.317 |  | 0.470 |  | 0.692 |  | 0.007 |  | 0.008 |  | < .001 |  | — |
| * p < .05, ** p < .01, *** p < .001 | | | | | | | | | | | | | | | | |

Supplementary Table 1: Spearman’s Correlations of probability of occurrence brain state and demographics before exercise.
