## Supplementary Table 2 for "Physical exercise and brain network dynamics: reduction of frontoparietal-striatal connectivity following 1 hour of aerobic cycling"

**Correlations post exercise**

| Spearman's Correlations | | | | | | | | | | | | | | | | | | | | | | | | | | | |
| --- | --- | --- | --- | --- | --- | --- | --- | --- | --- | --- | --- | --- | --- | --- | --- | --- | --- | --- | --- | --- | --- | --- | --- | --- | --- | --- | --- |
| **Variable** |  | | **1** | | | | **2** | | | **3** | | | **4** | | | **5** | | **6** | | **7** | | | **8** | | | **9** | |
| 1. Probability of occurrence brain state #1 post-exercise |  | Spearman's rho | |  | — |  | |  |  | |  |  | |  |  |  |  |  |  | |  |  | |  |  | |  |
|  |  | p-value | |  | — |  | |  |  | |  |  | |  |  |  |  |  |  | |  |  | |  |  | |  |
| 2. Probability of occurrence brain state #2 post-exercise |  | Spearman's rho | |  | -0.826 | *** | | — |  | |  |  | |  |  |  |  |  |  | |  |  | |  |  | |  |
|  |  | p-value | |  | < .001 |  | | — |  | |  |  | |  |  |  |  |  |  | |  |  | |  |  | |  |
| 3. Probability of occurrence brain state #3 post-exercise |  | Spearman's rho | |  | -0.730 | *** | | 0.409 | * | | — |  | |  |  |  |  |  |  | |  |  | |  |  | |  |
|  |  | p-value | |  | < .001 |  | | 0.047 |  | | — |  | |  |  |  |  |  |  | |  |  | |  |  | |  |
| 4. BMI |  | Spearman's rho | |  | 0.264 |  | | -0.190 |  | | -0.153 |  | | — |  |  |  |  |  | |  |  | |  |  | |  |
|  |  | p-value | |  | 0.212 |  | | 0.373 |  | | 0.474 |  | | — |  |  |  |  |  | |  |  | |  |  | |  |
| 5. Acitivity (min/week) |  | Spearman's rho | |  | 0.289 |  | | -0.276 |  | | -0.153 |  | | -0.232 |  | — |  |  |  | |  |  | |  |  | |  |
|  |  | p-value | |  | 0.170 |  | | 0.191 |  | | 0.474 |  | | 0.274 |  | — |  |  |  | |  |  | |  |  | |  |
| 6. VO2max (l/min) |  | Spearman's rho | |  | -0.027 |  | | -0.201 |  | | 0.244 |  | | 0.096 |  | 0.357 |  | — |  | |  |  | |  |  | |  |
|  |  | p-value | |  | 0.902 |  | | 0.346 |  | | 0.250 |  | | 0.655 |  | 0.087 |  | — |  | |  |  | |  |  | |  |
| 7. VO2max (ml/kg/min) |  | Spearman's rho | |  | -0.183 |  | | 0.010 |  | | 0.261 |  | | -0.539 | ** | 0.530 | ** | 0.667 | *** | | — |  | |  |  | |  |
|  |  | p-value | |  | 0.392 |  | | 0.963 |  | | 0.218 |  | | 0.007 |  | 0.008 |  | < .001 |  | | — |  | |  |  | |  |
| 8. Lactate pre-exercise |  | Spearman's rho | |  | -0.468 | * | | 0.299 |  | | 0.336 |  | | -0.389 |  | -0.259 |  | 0.075 |  | | 0.206 |  | | — |  | |  |
|  |  | p-value | |  | 0.021 |  | | 0.156 |  | | 0.108 |  | | 0.060 |  | 0.221 |  | 0.727 |  | | 0.333 |  | | — |  | |  |
| 9. Lactate post exercise |  | Spearman's rho | |  | -0.335 |  | | 0.252 |  | | 0.127 |  | | -0.236 |  | -0.286 |  | 0.060 |  | | -0.012 |  | | 0.412 | * | | — |
|  |  | p-value | |  | 0.110 |  | | 0.235 |  | | 0.556 |  | | 0.266 |  | 0.176 |  | 0.782 |  | | 0.956 |  | | 0.045 |  | | — |
| * p < .05, ** p < .01, *** p < .001 | | | | | | | | | | | | | | | | | | | | | | | | | | | |

Supplementary Table 2: Spearman’s Correlations of probability of occurrence brain state and demographics post-exercise.
